# Transcriptional Heterogeneity and Cell Cycle Regulation as Central Determinants of Primitive Endoderm Priming

**DOI:** 10.1101/2022.04.03.486894

**Authors:** Marta Perera, Silas B. Nissen, Martin Proks, Sara Pozzi, Rita S. Monteiro, Ala Trusina, Joshua M. Brickman

**Affiliations:** reNEW UCPH - The Novo Nordisk Foundation Center for Stem Cell Medicine, University of Copenhagen, Blegdamsvej 3B, DK-2200 Copenhagen N, Denmark; Niels Bohr Institute, University of Copenhagen, Blegdamsvej 17, DK-2100 Copenhagen Ø, Denmark; Department of Pathology, Stanford University School of Medicine, 291 Campus Drive, Stanford, CA 94305

## Abstract

During embryonic development cells acquire identity at the same time as they are proliferating, implying that an intrinsic facet of cell fate choice requires coupling lineage decisions to rates of cell division. How is the cell cycle regulated to promote or suppress heterogeneity and differentiation? We explore this question combining time lapse imaging with single cell RNA-seq in the contexts of self-renewal, priming and differentiation of embryonic stem cells (ESCs) towards the Primitive Endoderm lineage (PrE). Since ESCs are derived from the Inner Cell Mass of the mammalian blastocyst, ESCs in standard culture conditions are transcriptionally heterogeneous containing subfractions that are primed for either of the two ICM lineages, Epiblast and PrE. These subfractions represent dynamic states that can readily interconvert in culture, and the PrE subfraction is functionally primed for endoderm differentiation. Here we find that differential regulation of cell cycle can tip the balance between these primed populations, such that naïve ESC culture conditions promote Epiblast-like expansion and PrE differentiation stimulates the selective proliferation of PrE-primed cells. In endoderm differentiation, we find that this change is accompanied by a counter-intuitive increase in G1 length that also appears replicated *in vivo*. While FGF/ERK signalling is a known key regulator of ESCs and PrE differentiation, we find it is not just responsible for ESCs heterogeneity, but also cell cycle synchronisation, required for the inheritance of similar cell cycles between sisters and cousins. Taken together, our results point to a tight relationship between transcriptional heterogeneity and cell cycle regulation in the context of lineage priming, with primed cell populations providing a pool of flexible cell types that can be expanded in a lineage-specific fashion while allowing plasticity during early determination.

## Introduction

Naïve mouse Embryonic Stem Cells (mESCs) are karyotypically normal, immortal cell lines derived from the Inner Cell Mass (ICM) of pre-implantation embryos (Martin 1981; Evans and Kaufman 1981). ESCs are pluripotent, able to differentiate into all future cell lineages of the embryo proper, and able to retain pluripotency through successive rounds of self-renewing cell division (Beddington and Robertson 1989; Suemori et al. 1990; Lallemand and Brulet 1990; Martello and Smith 2014). In mouse, pluripotent stem cells can be derived from several stages of development and exhibit gene expression profiles matching these developmental stages (S. Morgani, Nichols, and Hadjantonakis 2017; Riveiro and Brickman 2020).

Under specific conditions, mESCs can recapitulate several aspects of the ICM specification. When cultured in serum and the cytokine LIF, they constitute a dynamically heterogeneous cell culture model that contains populations that are primed, but not committed, for both Primitive Endoderm (PrE) and Epiblast. These populations exhibit biases in differentiation but will readily interconvert when left in self-renewing culture (Canham et al. 2010; S. M. Morgani et al. 2013; Illingworth et al. 2016). While these conditions involve serum, we recently defined a culture condition that supports heterogeneous ESC culture, and at the same time sustains effective propagation of pluripotency and high levels of germline transmission (Anderson et al. 2017). We refer to this culture condition as NACL for N2B27, Activin A, Chiron and LIF. A similar defined media with the same components but a RPMI base (RACL) supports PrE differentiation and expansion (Anderson et al. 2017). As NACL ESC media is almost identical to the media used for PrE differentiation (RACL), this culture model is an ideal system for the comparison of self-renewal and differentiation.

Several time lapse studies have explored the role of heterogeneity in ESCs, and comparisons have been made between naïve culture in serum and media based on small molecule inhibitors of GSK3 and MEK, in addition to LIF (2i/LIF) (Singer et al. 2014; Abranches et al. 2014; A. Filipczyk et al. 2015; Cannon et al. 2015; Hastreiter et al. 2018). Taken together, these studies have focused on the dynamic heterogeneity of pluripotency factors which are associated with supporting the pluripotent state and Epiblast specification *in vivo*. Although they have explored the role of pluripotency factors in lineage priming (Strebinger et al. 2019), they have not assessed the expression of lineage-specific markers, how cells progress into differentiation and the role of priming in differentiation. Here we explore these questions with respect to PrE priming and differentiation.

*In vivo*, the segregation of PrE from Epiblast is regulated by the FGF/ERK pathway (Chazaud et al. 2006). The inhibition of this pathway results in an expansion of Epiblast identity and loss of PrE, both *in vivo* and *in vitro* (Nichols et al. 2009; Yamanaka, Lanner, and Rossant 2010; Hamilton and Brickman 2014; Saiz et al. 2016). Based on the activity of this pathway in supporting PrE differentiation *in vivo*, inhibition of the kinase upstream ERK, MEK, with a pharmacological antagonist (PD03) is an important component of the defined ESC culture system known as 2i/LIF (Ying et al. 2008). Additionally, ERK activation is also associated with cell cycle progression and is thought to stimulate G1/S transition (Yamamoto et al. 2006; ter Huurne et al. 2017). Although G1 lengthening has been functionally related to endoderm differentiation (Calder et al. 2013; Pauklin and Vallier 2013; Coronado et al. 2013), it is not clear how ERK stimulation of cell cycle progression relates to its capacity to induce differentiation.

Here we focus on the dynamics of PrE priming and differentiation exploiting both time lapse imaging and single cell RNA-seq to link both events to regulation of the cell cycle. ESC culture (NACL) was found to support the more rapid proliferation of the Epiblast-primed population, and PrE differentiation (RACL) promotes the more rapid proliferation of cells primed for endoderm differentiation. We found that FGF signalling regulates cell states and proportions through both coordinating inheritance of cell cycle lengths as well as rates of endoderm priming. This functional regulation of cell cycle in heterogeneous cell cultures indicates that cell cycle synchronisation supports self-renewal as well as preparing cells for further differentiation. Furthermore, we found that G1 length is adjusted independently from the cell cycle, pointing to a model where cells receive signals from cytokines during G1, while still actively proliferating. Taken together, our work suggests that cell cycle length is not only tightly regulated by the culture context, but that this coordination has a functional role in both heterogeneity and lineage choice.

## Results

### Single cell RNA-seq of PrE differentiation reveals the presence of both endoderm and Epiblast-like populations in differentiation

To assess lineage-specific transcriptional heterogeneity in PrE differentiation and compare the events occurring *in vitro* to those occurring *in vivo*, we performed single cell RNA sequencing by MARS-seq (Jaitin et al. 2014) on samples from 2i/LIF and days 1, 2, 3, 4 and 7 of *in vitro* PrE differentiation (Fig 1A). To track both endodermal and Epiblast lineages, we used a Sox2-GFP and Hhex-mCherry double fluorescent reporter cell line (Fig 1A) (Anderson et al. 2017). Indexing based on reporter expression and plate-based MARS-seq enabled us to link individual transcriptomes to the cell types identified by FACS (Fig S1A-B). We collected equivalent numbers of different populations (Sox2 High and Low) to ensure we had a good representation of the spectrum of cell types occurring in these *in vitro* cultures. As a result, the cell proportions based on cluster composition do not reflect the proportion of these cell types present at different time points during differentiation.

**Figure 1.**
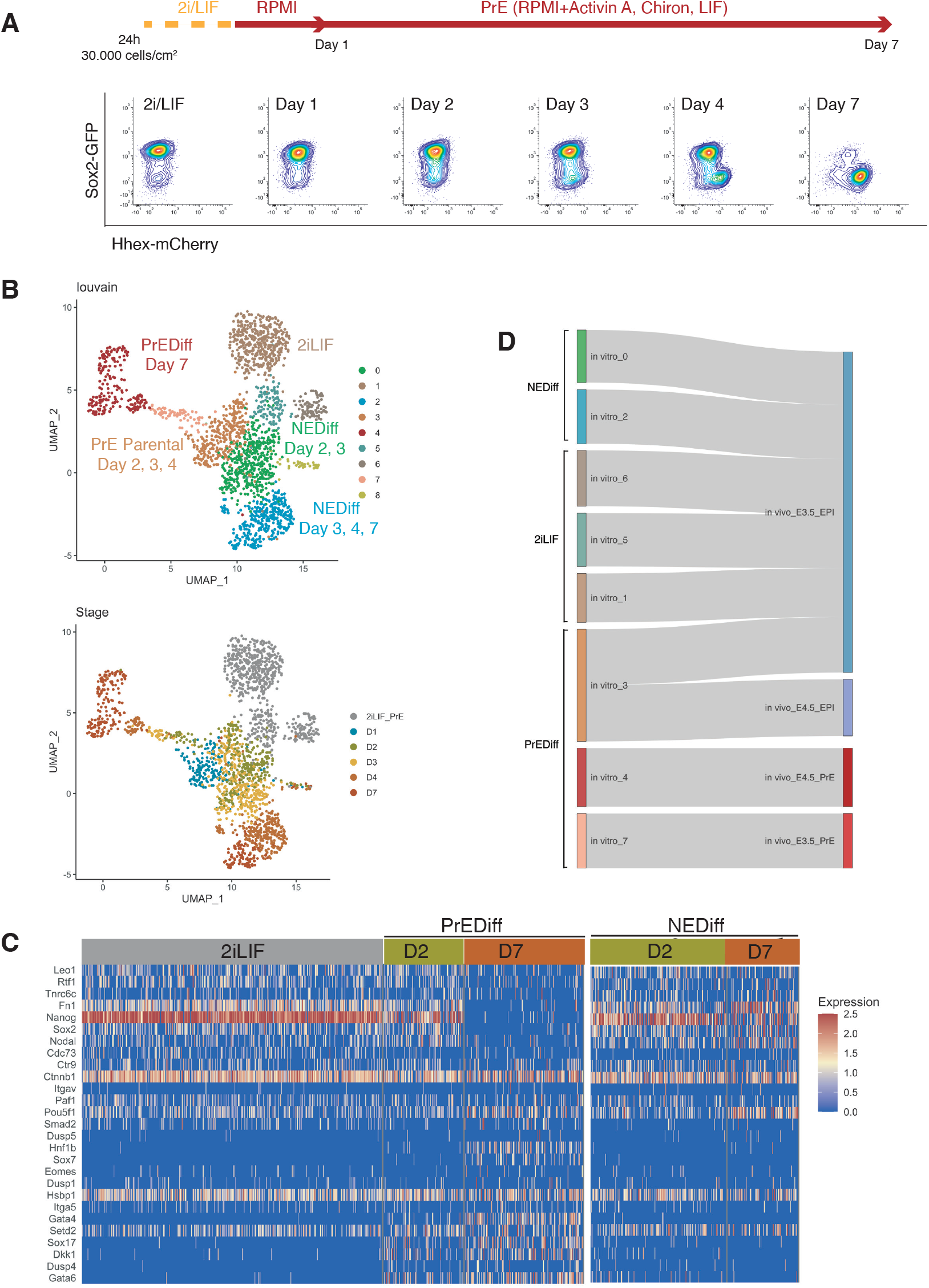
Transcriptome profiling of PrE *in vitro* differentiation. A. Schematic of the experiment. Cells were passaged twice in 2i/LIF and then plated in RPMI base media 24 hours before starting the experiment. Bottom panel: Flow cytometry plots showing the time points selected for single cell RNA-seq. The fluorescent information of Sox2 and Hhex was recorded prior to sequencing. B. UMAP projection of the *in vitro* experiment showing 9 identified clusters using Louvain (upper panel) and stages of differentiation (bottom panel). C. Heatmap of endoderm genes (GO term 0035987) expression in 2i/LIF, day 2 and day 7 of differentiation. Left panel: PrE Diff branch. Right panel: NEDiff branch. Cells at day 2 in the PrE branch already are upregulating endoderm genes while the NEDiff cells are not. D. Sankey plot visualizing cluster similarity comparison between identified *in vitro* clusters and *in vivo* (Nowotschin et al. 2019) experiment using in-house Cluster Alignment Tool (CAT).

We detected 9 clusters using unsupervised clustering (Fig 1B). Clusters 1, 5 and 6 were composed of 2i/LIF cells (Fig S1C, Table S1). Differentiating cells (from days 2 to 7) appeared to resolve into two branches: a PrE-like branch (clusters 3, 7 and 4) expressing progressively increasing levels of Hhex-mCherry (Fig S1A) and PrE markers such as Dab2, Gata6, Pdgfra, and Sox17 (Fig S2A); and an Epiblast-like branch (clusters 0 and 2) with cells that appear to remain Epiblast-like, continuing to express pluripotency markers (Sox2-GFP, see FigS1B. Klf2, Nanog, Sox2, and Rex1, see Fig S2B). As most Epiblast-like cells maintained expression of pluripotency markers (Fig. S2B), but clearly have distinguished themselves from 2i/LIF ESCs, we refer to these cells as Non-Endodermal/Non-Differentiated (NEDiff) (Fig 1B).

We identified distinct endodermal signatures as early as day 2 (Fig 1C), suggesting that some cells at day 2 were already primed towards endoderm differentiation. The endodermal genes found at day 2 of the PrE branch (PrEDiff in Fig 1C) were not upregulated at day 2 in the NEDiff branch (NEDiff in Fig 1C), supporting the notion that the transcriptional signature for PrE appeared on day 2 in a sub-population of these cultures.

To compare these cell types to those induced *in vivo*, we compared our dataset with the already published scRNA-seq dataset of pre-implantation embryos (Nowotschin et al. 2019) using the Cluster Alignment Tool (CAT) (Rothová et al. 2022). Our day 7 PrE cluster 4 was most similar to PrE from E4.5, the earlier emerging PrE cluster 7, first appearing at day 2, resembled PrE from E3.5, while the rest of the clusters were most similar to E3.5 Epiblast (Fig 1D). Although cluster 3 was positioned at the beginning of the PrE branch, it aligned to Epiblast, continued to express Sox2 and showed low levels of Hhex. Based on these comparisons, we believe that our *in vitro* model is a good tool for deconstructing differentiation and that an early PrE-like cell type arises by day 2 *in vitro*.

### PrE arises through early induction, followed by selective proliferation

We next sought to determine whether we could detect the emergence of these early (day 2) PrE-like cells and follow their differentiation using time lapse microscopy. We exploited a mESC line that couples a sensitive PrE reporter, Hhex-mCherry, with a second that enables lineage tracing, H2B-Venus (HFHCV) (Illingworth et al. 2016). These cells were used to follow mESC in defined PrE differentiation (RACL) (Fig 2A). To follow a significant number of cell cycles through differentiation, we acquired images every 20 minutes for 6 days (See Supplementary Movie 1).

**Figure 2.**
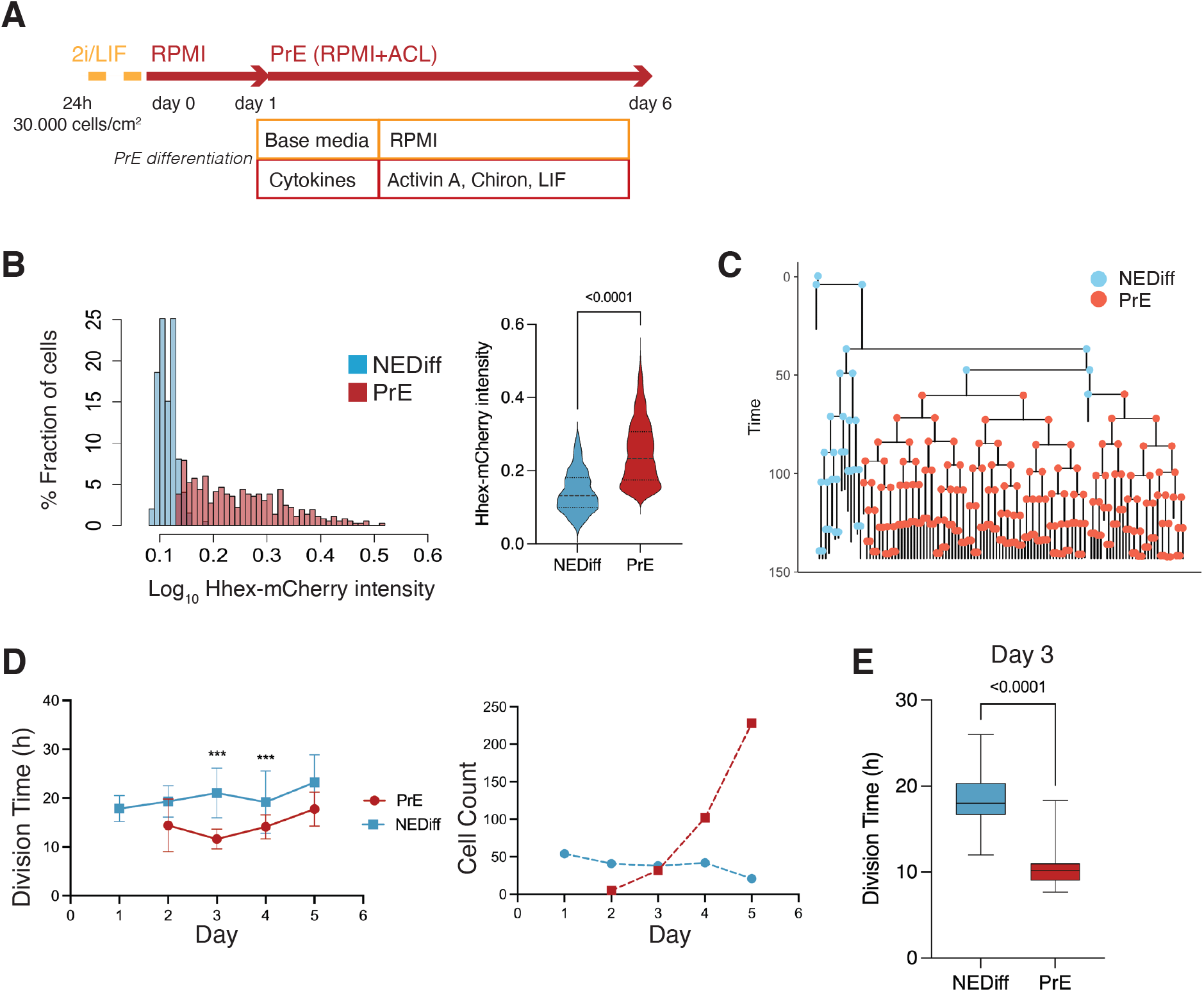
Time lapse of PrE differentiation shows rapid proliferation of PrE primed cells. A. Schematic of the experimental setup. Cells were imaged for 6 days acquiring 1 time frame every 20 minutes. B. The *Hhex* intensity distribution between the populations allowed us to separate PrE differentiated cells (PrE) from the Non-endodermal differentiated cells (NEDiff). p-value < 0.0001 Mann-Whitney test. C. Example of a lineage tree showing how the PrE branch of the tree arises. The first and last generation were discarded from further analysis since the cell cycle information is not completed. All lineage trees collected in the PrE condition are shown in Figure S3; this example corresponds to Tree 10 in Figure S3. D. Analysis of mESCs division times and cell counts showed that cells that differentiate into PrE are dividing faster at the beginning of the differentiation process (day 3, left panel) and that selected survival takes place in the last days of differentiation (day 4 and 5, right panel). *** p-value < 0.001. E. Cell cycle length at day 3 shows fastening in the PrE cells division time, compared to a slower dividing non-endodermal cluster (p-value<0.0001 Unpaired t-test).

As with the scRNA-seq time course, we started differentiation from a relatively uniform population of cells cultured in 2i/LIF and then differentiated them to PrE (See Methods, Fig 2A). When we analysed the fluorescence intensity distribution in this setup, we observed a distribution that suggested these cultures contained the same two populations as observed in the scRNA-seq: differentiated PrE and the NEDiff population that failed to progress towards endoderm. We clustered cell intensities using k-means clustering and used this clustering to assign identity in differentiation (Fig 2B, S3). The identity and distribution of PrE and NEDiff states were confirmed with GATA6 and NANOG staining (Fig S4A). Based on the lineage trees (Fig 2C and S3) and increasing levels of PDGFRA expression (Fig S4B), the endodermal identity appeared around day 3 from a small founder population. This was also apparent when viewing individual time lapse movies (Supplementary Movie 1).

As possible contributors to the growing PrE population across *in vitro* differentiation, we aimed to distinguish between a selective or progressively inductive process. Selection would imply faster proliferation or enhanced survival of a founder population of primed PrE cells, while induction would be represented by endoderm cells converting continuously from the undifferentiated cell pool. To evaluate these possibilities, we assessed the behaviour of both endodermal and non-endodermal populations over time. We observed a decrease in the division time of the PrE cluster with cell division accelerating primarily on day 3 of differentiation (13.00 ± 4 h for PrE and 20.00 ± 6 h for NEDiff clusters on day 3) (Fig 2D-E, Table S2). This is the point in time when PrE clones became distinguishable from NEDiff cells in the time lapse video (Supplementary Movie 1) and suggested that differentiation proceeded, in part, by providing an environment that favoured the proliferation of a PrE-primed population.

As these cultures started from a homogeneous population of 2i/LIF cells, we assumed some conversion to endoderm fate preceded the acceleration of proliferation and assessed the probability of cell state transition before and after day 3 (72h). We found that 12.6%± 0.6 of the total population upregulated *Hhex* expression during the first 72h, thereby transitioning into the PrE cluster. However, after the 72h time point, only 1.9% ± 1.2 of the cells were able to transit between clusters. This suggested that competence to enter the endoderm lineage was lost after the first 3 days of the differentiation, indicating that proliferation was enhanced following lineage choice. Hence, even though only a small population of cells entered differentiation, the selective expansion of these cells allowed them to take over the final culture.

To assess whether the combination of this small amount of lineage conversion followed by enhanced proliferation was sufficient to account for the behaviour of differentiating cultures, we sought to generate a simple model introducing the transition probability observed in time lapse (*t* in Fig 3A). This minimal set of parameters was not enough to recapitulate our lineage trees. We found that only after introducing differential survival rates for PrE and NEDiff cells (*s* in Fig 3A) we could reproduce the cell populations observed. Thus, this minimal model predicted that selective cell death was also required to explain differentiation. To quantify the extent of this during differentiation, we determined the survival rates in our cultures. Before commitment took place at day 3, 86% of prospective PrE cells survived to generate colonies of PrE, whereas only 61% of the NEDiff survived (Table S2). This difference in survival was further enhanced following commitment at 72 hours: from this point onwards, 92% of cells in the PrE cluster survived, whereas only 39% of the cells in the NEDiff cluster did (Table S2). Therefore, it appeared that differentiation was selecting for committed endoderm at two levels: proliferation and cell survival. Furthermore scRNA-seq suggested this population also accrued more endodermal identity as it expanded.

**Figure 3.**
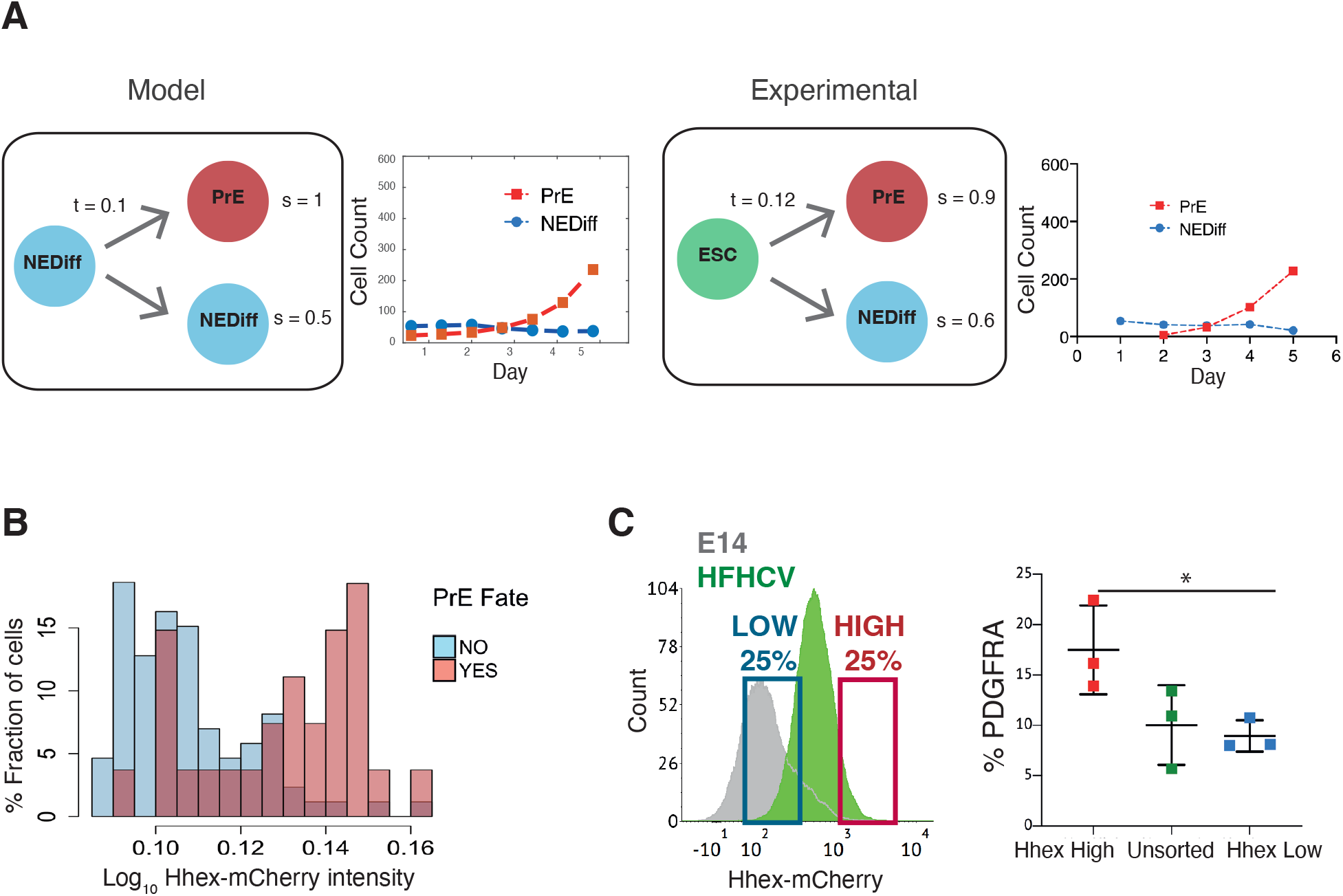
Analysis of PrE progenitor cells demonstrates functional priming of Hhex-high cells. A. A mathematical model that only considers the difference in cell death rate between the 2 populations can recapitulate the PrE dataset collected (see Methods for description of the Mathematical Modelling). Looking into the experimental dataset, we found the same survival rates that the model predicted. t = transition rate, s= survival rate. B. The NEDiff cluster at day 2 was separated into cells that will give rise to PrE (PrE Fate YES), and cells that eventually would not differentiate (PrE Fate NO). Analysis of the *Hhex* intensity distribution shows that cells that will give rise to PrE (PrE Fate YES) show higher *Hhex* intensity. Total cell number and fluorescence quantification shown in Table S3. C. The High Hhex population from day 2 of PrE differentiation, isolated by FACS, shows improved PrE differentiation (scored as percentage of PDGFRA positive cells), demonstrating the functional priming of these cells. * p-value < 0.05 Kruskal-Wallis test.

Lastly, given that we found an endodermal transcriptional signature in the scRNA-seq at day 2, we considered whether lineage priming and selection could be occurring prior to our assignment of differentiation (day 3). Thus, we took the dataset from day 2 of differentiation and asked whether low level reporter expression correlated with later differentiation by asking which cells would prospectively give rise to PrE (PrE Fate) and which would never become endoderm by day 3 (Not PrE Fate). We found that there was a tendency for PrE progenitor cells (PrE Fate) to express *Hhex* at higher levels (0.18) compared to cells that are non-endodermal progenitors (Not PrE Fate, 0.15) (Fig 3B, Table S3). To confirm the notion that these cells were primed functionally at day 2, we isolated *Hhex*-high and low populations from day 2 of the PrE differentiation by FACS and cultured them further in PrE differentiation conditions. Consistent with the time lapse analysis, we observed that the day 2 Hhex-high cells produced significantly more PDGFRA positive PrE than either the Hhex-low or unsorted populations (Fig 3C).

### PrE-primed mESCs in steady state culture conditions

As the priming observed during *in vitro* PrE differentiation was similar to the spontaneous dynamic priming that occurs in standard ESC culture, but primed PrE remained a minority of the culture, we hypothesized that ESC culture might favour the proliferation of the undifferentiated or Epiblast-like population. As ESC culture can be supported by the same cytokines but with a different base media (Anderson et al. 2017), we reasoned the change in base media might underlie this difference. To explore this issue, we performed time lapse microscopy on the HFHCV mESC line in chemically defined NACL culture (Anderson et al. 2017) (Fig 4A). We observed expansion of mESCs colonies from a small set of 2-8 cells into several hundreds of cells, which produced imaging of up to 7 generations (See Supplementary Movie 2, Fig 4B).

**Figure 4.**
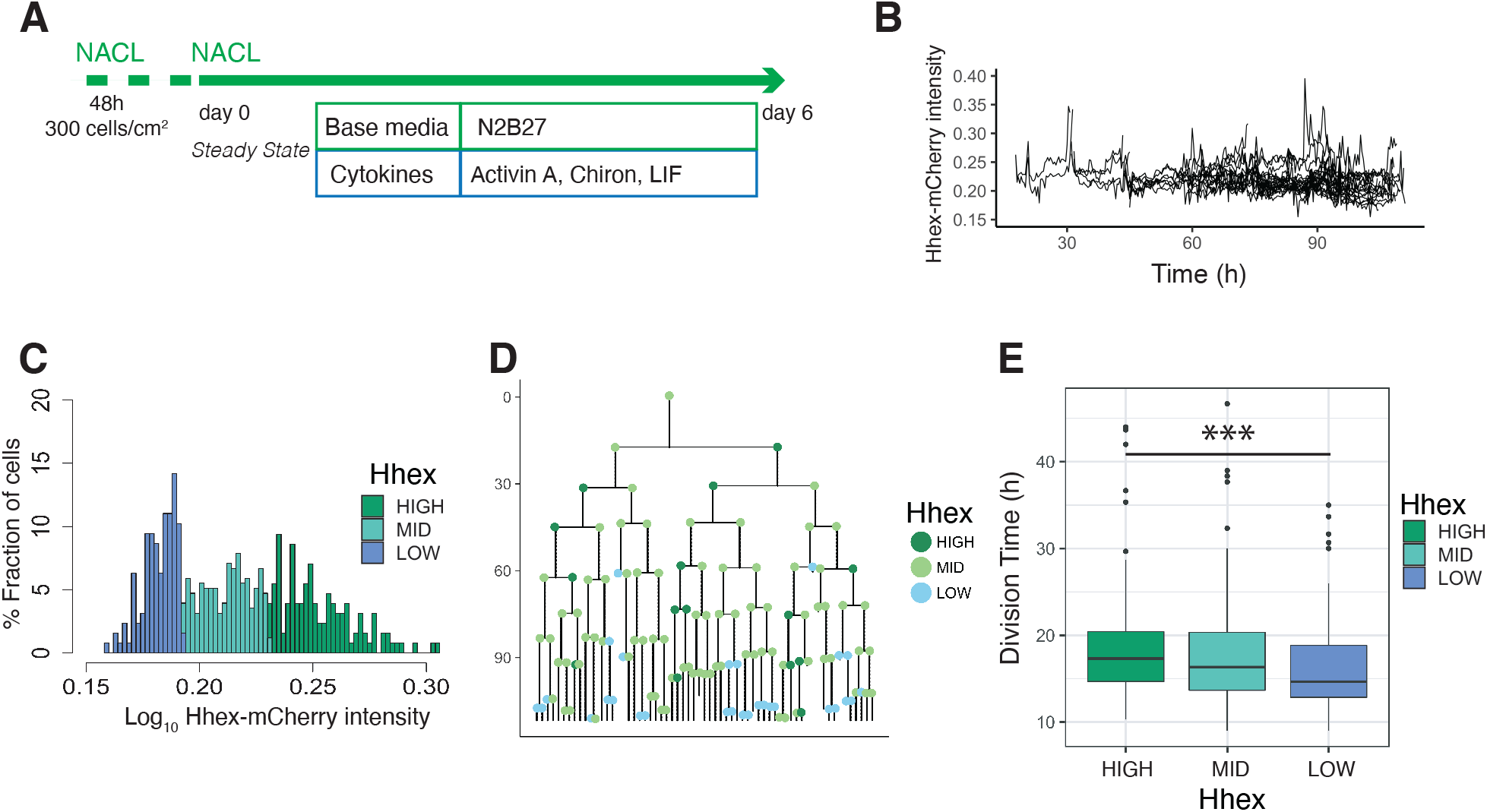
Single cell quantification of *Hhex* expression in NACL uncovers a relationship between *Hhex* levels and cell cycle length. A. Schematic of the experimental setup. Cells were plated 48 hours before starting the experiment. Cells were imaged for 6 days acquiring 1 time frame every 20 minutes. B. Example of a cell trace (Time vs Hhex-mCherry intensity) in the setup analysed. Cells survived and divided over 6 days without any apparent effect of cell death. Cells were entering and exiting higher and lower *Hhex* states without any apparent bias. C. *Hhex* intensity distribution was divided into 3 compartments: High (includes cells above 75% percentile), Mid (between 25% and 75% percentiles) and Low (cells below 25% percentile). Y-axis shows the percentage of cells that falls into each bin. See Table S4 for total cell numbers per compartment. D. Example of a lineage tree with the corresponding compartments of *Hhex*, by colour. The first and last generation were discarded from further analysis since the cell cycle information is not completed. All lineage trees collected in the NACL condition are in Figure S6. This example corresponds to Tree 1 in Figure S6. E. The Low Hhex population divides significantly faster than the High Hhex. *** p-value <0.001, Kruskal-Wallis test. The Mid Hhex population shows an in-between division time, suggesting a linear relationship between *Hhex* expression level and cell cycle length.

We confirmed our previous observations (Anderson et al. 2017) that NACL culture supports a population of ESCs primed for PrE (Hhex-mCherry positive) and Epiblast (NANOG positive), within the OCT4 positive ESC population (Fig S5A). The distribution of *Hhex* expression that corresponded to these states was quite broad and showed a non-normal distribution (Fig S5B), similar to that observed for the constitutive lineage label H2B-Venus. Ideally, if the PrE-like and Epiblast-like states were well defined, one would expect a bimodal distribution with one peak for each state. However, similar to previous reports for fluorescent reporters in ESC culture (Abranches et al. 2014; Singer et al. 2014), this was not observed. Based on previous analysis of NANOG expression (A. Filipczyk et al. 2015; Hastreiter et al. 2018), we divided the intensity distribution into compartments based on quartiles of the mean fluorescence intensity measurements. The compartments were called High (cells within the highest 25% intensities), Low (cells within the lowest 25% intensities), and Mid (the rest of the intensities) (Fig 4C). We constructed lineage trees using this compartmental distribution (Fig 4D, Fig S6).

The intersection between selection and induction during PrE differentiation time lapse described how a primed progenitor pool could give rise to a differentiated cell population. In NACL culture, primed cells are heterogeneously maintained, but the mechanisms that support and restrain this population are unknown. Therefore, we sought to determine how cell death, transitions, and cell cycle sustain naïve and primed populations in a dynamic equilibrium at steady state.

In NACL culture, we observed that 86% of cells survived, and cells divided every 17h on average. The level of cell death was not significantly different between the three compartments in our dataset (9% in the High, 12% in the Mid, 21% in the Low, Table S4). Based on the lineage trees (Fig S6), we estimated the probability that a cell would transit between these states by assuming that a cell undergoes transition if its mean intensity, averaged across cell cycle, changes the compartment. Altogether, we observed a similar behaviour between the Mid compartment and either the High or Low compartment (Fig S5C). In RACL (PrE media) the Hhex-expressing primed population had a short cell cycle, but in NACL, where ESC self-renewal predominates, we found the opposite. Here, the Low Hhex cells exhibited significantly shorter division times than the High Hhex cells (14.67± 5 h vs 17.33 ± 6 h, Fig 4E, Table S4). This result suggests a relationship between lineage priming and cell cycle length, with pluripotency being supported by media conditions that promote faster cycling of Epiblast-primed cells.

### MEK inhibition affects mESC priming by increasing the fraction of fast proliferating cells

While transition probabilities favour the expansion of the Mid compartment, shorter cell cycle times favour the Low compartment, and cell death appears indiscriminate. As FGF/ERK signalling both regulates PrE specification by directly altering enhancers and coordinates the cell cycle promoting G1/S and G2/M transitions (Yamamoto et al. 2006; Hamilton and Brickman 2014) we asked whether inhibiting this pathway with PD03 acts to accelerate cell state transitions, cell cycle or both. We quantified the changes in both *Hex* expression and cell cycle in response to FGF/ERK inhibition by PD03 (Fig 5A, Fig S7), and the 3 compartments were defined using the same threshold intensities as for the NACL dataset.

**Figure 5.**
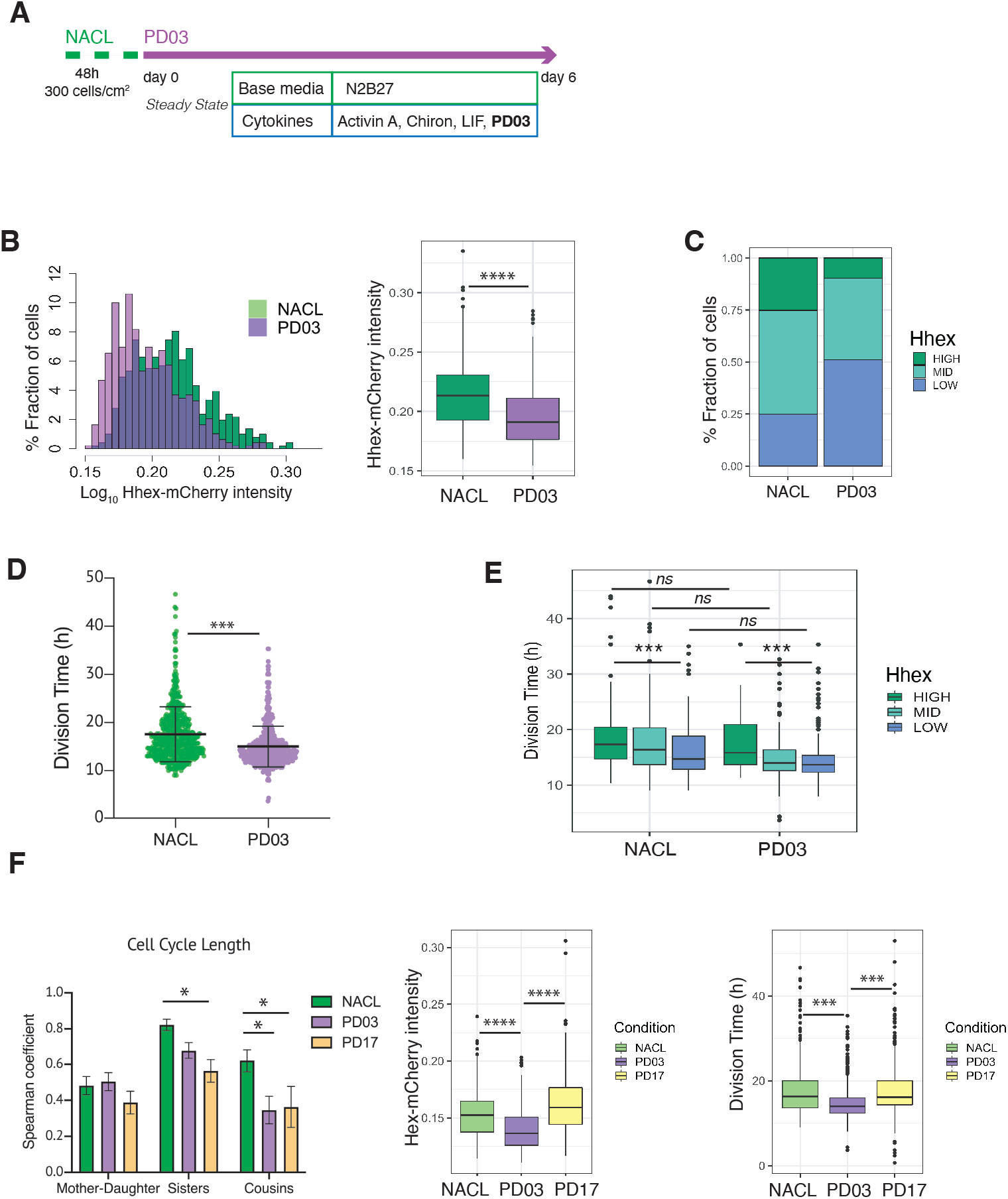
PD03 promotes expansion of the fast proliferating Low Hhex population. A. Schematic of experimental setup. Cells were plated 48 hours before starting the experiment in NACL, and PD03 was added at the start of the time lapse. Cells were imaged for 6 days acquiring 1 time frame every 20 minutes. B. *Hhex* intensity is significantly lower in the PD03 treated population. **** p-value < 0.0001, Wilcoxon test. C. The lower intensity of *Hhex* is related to the higher fraction of cells in the Low Hhex population. The Low Hhex population increases from 25 to 50% when PD03 is added. D. mESCs division time is significantly faster in PD03. *** p-value < 0.001, Mann-Whitney test. E. Cell cycle in the Low Hhex compartment is faster in PD03 as well as in NACL. *** p-value < 0.001, Kruskal-Wallis test. F. Left: Both PD03 and PD17 produce a loss in the cell cycle synchronisation between sisters and cousins. All correlation plots are shown in Figure S8. * p-value < 0.05. Right: PD17 does not provide the same alterations in *Hhex* expression or division time that were generated by PD03 *** p-value < 0.001, **** p-value < 0.0001.

Given the direct relation between ERK stimulation and *Hhex* upregulation (Hamilton and Brickman 2014), we found unsurprisingly that cells in PD03 expressed significantly lower levels of *Hhex* than in the NACL control (Fig 5B). Thus, the proportion of cells in the Low Hhex compartment increased (Fig 5C). To understand if this increase stems from changes in cell death, proliferation or switching probabilities, we compared these between the NACL and PD03 conditions. At the level of cell survival, PD03 treatment had little effect (80% cells that survived vs 86% in NACL, Table S4). However, culture in PD03 produced significant alterations to both cell cycle length (Fig. 5D) and cell state transitions (Fig. S8A). While the influence of PD03 on cell state transitions is difficult to quantify robustly as PD03 can induce rapid cell state changes (Hamilton et al. 2019), we observed a high transition rate from the rare High Hhex cells found in PD03 conditions. This suggests that inclusion of PD03 in cell culture media suppresses the stability of the High state, such that any cell that escapes the Low Hhex and Mid compartments, rapidly returns.

We observed a shorter division time in the population of cells treated with PD03 (Fig 5D and Table S4, confirmed by bootstrap analysis, Fig S8B). Similar to the results observed in NACL, in PD03, cells in the Low Hhex compartment (Epiblast-like) had a shorter cell cycle than those in the other two compartments (Fig 5E, Table S4). However, when we compared the division time in the same compartment, but different media conditions (i.e., plus and minus PD03), we did not observe significant differences (Fig 5E). Together with an increased fraction of cells in the Low Hhex compartment (Fig 5C), this indicated that on a population level, PD03 decreased cell cycle length by increasing the fraction of fast proliferating cells in Low Hhex (Epiblast-like) state. Thus, the change in cell cycle length observed in Figure 5D appears to be a population effect and not a result of PD03 regulating the cell cycle at the level of individual cells.

It has previously been shown that cell cycle lengths are inherited. This does not occur transgenerationally, as mothers and daughters do not have correlated cell cycle lengths, but sisters and cousins do (Sandler et al. 2015). Does this imply that lineage priming is heritable as well? In NACL, we found that cell cycle length was correlated between sisters and cousins, but not mother-daughter (Fig S8C). Given that PD03 produced a more homogeneous population of Hhex Low cells that divided faster, we hypothesized that cell cycle inheritance might therefore be better correlated. However, we found that cell cycle correlation between sisters and cousins was reduced or eliminated in the presence of PD03 (Fig 5F, Fig S8D). This capacity of PD03 to inhibit cell cycle correlations suggests that the mechanisms governing cell cycle entrainment are likely dependent on signalling downstream of MEK.

We considered two possible paradigms for ERK-dependent regulation of cell cycle synchronisation: either the symmetric inheritance of a kinase activity, or the paracrine activity of a cytokine that drives MEK/ERK activation. FGF4 is a prominent cytokine known to activate the MAPK/ERK in ESCs and early embryos (Kunath et al. 2007; Yamanaka, Lanner, and Rossant 2010). Moreover, limited durations of FGF/ERK signalling can induce ESCs to adopt a PrE primed state homogenously (Hamilton and Brickman 2014). This suggests that the paracrine signalling with FGF and its receptor (FGFR) may be responsible for cell cycle synchronisation in ESC culture. To test this idea, we followed ESC cell cycle and lineage heterogeneity in response to pharmacological inhibition of FGFR with PD17 (PD173074, an FGFR1/3 inhibitor). Figure 5F shows that culture in NACL with PD17 resulted in the same loss of cousin and sister correlation as that in response to inhibition of its downstream kinase MEK, further supporting the role for this pathway as a primary determinant of cell cycle synchronisation. Furthermore, PD03 was both more effective at inducing Low Hhex populations and producing a general decrease in cell cycle length (Fig 5F right; Table S4). However, despite the reduced effect of PD17 in ESC heterogeneity (Fig 5F right), PD17 was at least as effective as PD03 in its capacity to block synchronisation (Fig 5F left; Fig S8E). Thus, even though PD17 containing cultures had significant proportions of Mid Hhex and High cells, that proliferated slowly, PD17 still blocked their synchronisation, suggesting that this pivotal pathway independently regulates cell state transitions and cell cycle synchronisation.

### G1 length regulation accompanies the changes observed in cell cycle

To further elucidate the differences in cell cycle length in this context, we performed time lapse using the FUCCI cell cycle reporter cell line (Sakaue-Sawano et al. 2008), engineered with an H2B lineage reporter (See Methods, Supplementary Movie 3). Using this approach, we measured division time, G1 length and G1 in relation to the total cell cycle length (G1 Ratio) (Fig 6, Table S5). We measured these parameters in NACL, PD03, 2i/L and PrE differentiation.

**Figure 6.**
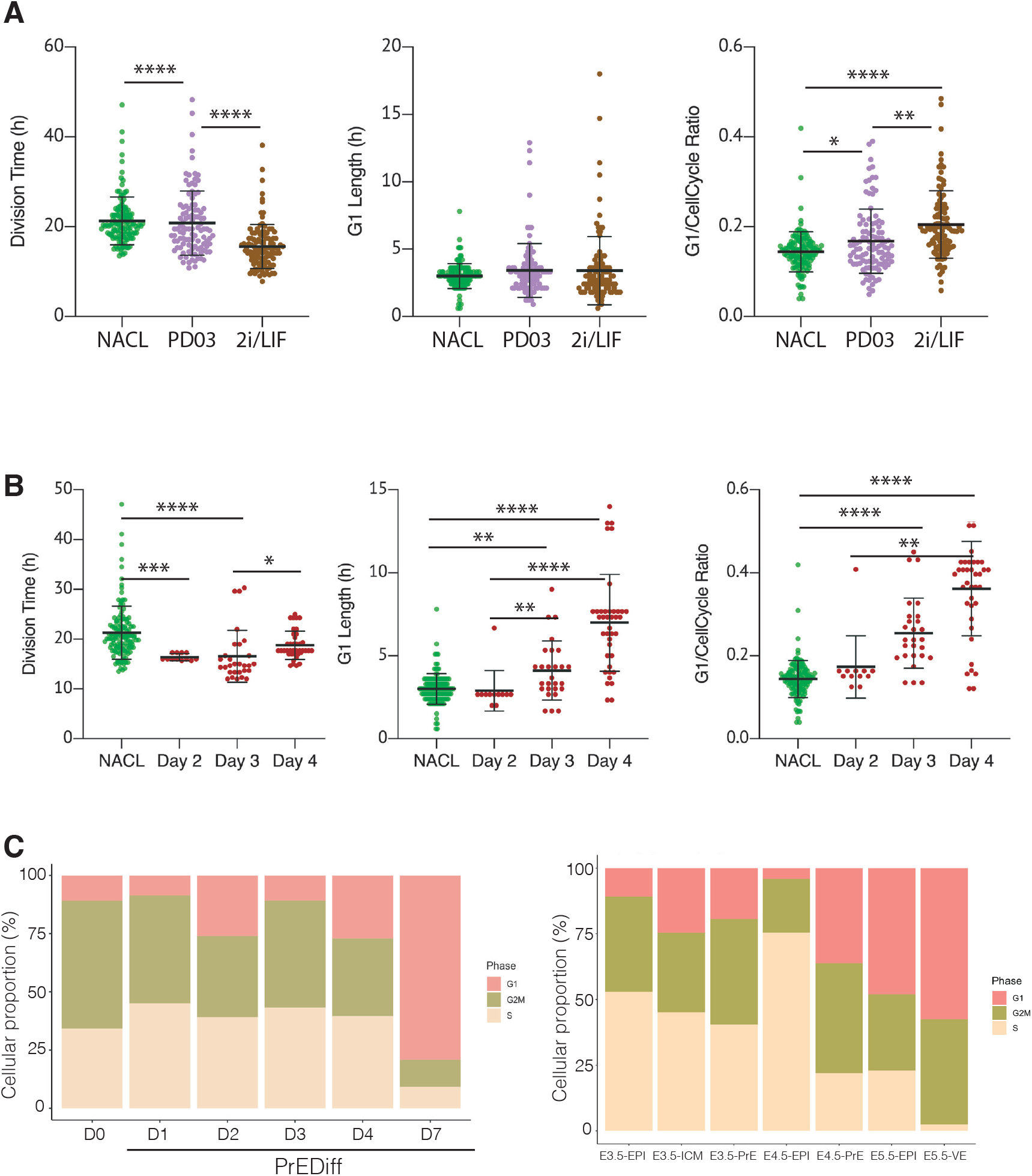
The regulation of cell cycle length during priming and differentiation involves G1 length. A. PD03 addition significantly increases the ratio between G1 and Division Time (* p-value < 0.05, Kruskal-Wallis test), even though it does not change significantly the G1 length in FUCCI cells. Cells cultured in 2i/LIF for 2 passages show a faster cell cycle than when PD03 was added for a short period (** p-value < 0.01, **** p-value < 0.0001 Kruskal-Wallis test). B. PrE differentiated cells show a longer G1 phase and an increase in G1 Ratio as they go along the differentiation process. Division time is faster at day 3 and then it slows down, consistent with the previous dataset. * p-value < 0.05, ** p-value < 0.01, *** p-value < 0.001, **** p-value < 0.0001 Kruskal-Wallis test. C. Proportion of cells that express G1, G2/M or S signature transcriptional profiles. Left: *in vitro* dataset (this study). Right: *in vivo* dataset (Nowotschin et al. 2019).

Comparing NACL, PD03 and 2i/L, PD03 did not seem to affect the average G1 length significantly, yet there appeared a small increase and clear outliers within PD03 and 2i/LIF cultured cells that showed an increased G1. Although the significance of this increased G1 length may require additional measurement, the increase in G1 length relative to cell cycle length was significant. Thus, G1 either remains unaffected or is marginally increased despite the decreasing cell cycle time (Fig 6A, Table S5). We confirmed these unexpected changes by bootstrap analysis (Fig S9). Collectively our results suggest that G1 length is increased relative to the cell cycle in response to PD03. As we cannot combine the G1 reporter with *Hhex* lineage, we cannot assess whether it is the High Hhex or Low Hhex cells which present a longer G1. However, as the impact of PD03 on cell cycle is a consequence of stimulating the Low Hhex population, we presume an increase in this population accounts for the change in the ratio of G1 to the cell cycle. Taken together, our results suggest that PD03 promotes the faster proliferating Low Hhex population, but that the decreased cell cycle time in this population is not a result of stimulating G1/S transition, consistent with the role of this drug in inhibiting it (ter Huurne et al. 2017).

We also assessed alternation in G1 during PrE differentiation (See Supplementary Movie 4). Since this cell line did not have a *Hhex* marker, we stained for GATA6 at the end of the time lapse and retrospectively identified PrE lineage trees, mapping the progenitor cells that would give rise to GATA6 positive colonies. Using this approach, we were able to locate the progenitors of fully differentiated PrE cells and determine their cell cycle parameters on different days of differentiation (Fig 6B; Table S5). Despite the decreased cell cycle length observed for endoderm committed cells at day 3, we observed a progressive increase in G1 itself and the G1 Ratio (Fig 6B, right). This increase in G1 appeared in parallel with the increase in proliferative capacity that we assumed associated with endoderm commitment arising from day 3 to 4, further strengthening the apparently contradictory correlation between increased G1 and higher rates of cell division. While there is an enhanced ratio of G1 length in Epiblast-primed ESCs, the changes in G1 during differentiation are much more significant, with G1 itself doubling as cells progress in differentiation.

Finally, we assessed the proportion of cells that expressed G1 transcripts on our scRNA-seq dataset from *in vitro* differentiation. As shown in Fig 6C, we observed the same trend of cells entering G1 as they progress through the endoderm lineage. Taking advantage of the published early embryo dataset used to benchmark our differentiation (Nowotschin et al. 2019), we assessed whether a G1 trend also occurs in vivo during PrE specification. We found an increase in cells expressing G1 transcripts as they progressed into the endodermal lineage *in vivo*. Around the time of implantation E4.5, there was a robust difference in cell cycle phase between Epiblast, which was mostly in S phase, and PrE, where almost half of the cells were in G1, suggesting the coupling we observed between cell cycle and differentiation *in vitro* may be recapitulated in endoderm specification *in vivo*.

## Discussion

In this paper we describe a close relationship between cell cycle length, lineage priming and differentiation, both *in vivo* and *in vitro*. In ESC self-renewal *in vitro*, naïve Epiblast-like cells proliferate faster than those primed for PrE differentiation, but when the base media is changed, the tides are turned, and the PrE-primed population develops the proliferative advantage. This suggests that differential culture conditions identify the appropriate lineage-biased populations and stimulate their relative expansion. Our observation that similar stages exist in peri-implantation development suggests that while this facet of differentiation is exploited *in vitro*, it is also a fundamental component of cell choice *in vivo*.

The differentiation process we define is both selective and progressively inductive. Heterogeneities arise, and these appear to be captured in early differentiation. However, the lineage-specific nature of these primed cells increases inductively. Primed cells that fail to enhance endodermal gene expression tend to die. By day 3 in differentiation, all the future founder cells in the culture are set, and further commitment does not occur. At this point, a combination of selective cell survival, increased rates of proliferation and enhanced endodermal transcription lead to the final differentiated cell populations. However, while these conditions can promote more rapid cell cycling, they produce apparently contradictory increases in relative G1 lengths.

What are the factors that govern a cell’s choice to differentiate? An increase in G1 would give cycling cells more time to respond to inductive signals, ultimately leading to increased responsiveness to differentiation cues. This would suggest that cells progress into early differentiation, lengthening their G1 phase, thereby improving accessibility to cues promoting commitment, and enabling signalling dependent transcription factors to accumulate ahead of replication. While the link between G1 phase and commitment of pluripotent stem cells into differentiation was first described in 1987 in embryonal carcinoma cells (Mummery, van den Brink, and de Laat 1987), the recent development of the cell-cycle reporter cell line FUCCI has allowed for more detailed findings in this field. It has been functionally demonstrated that the G1 fraction of ES cells has an enhanced capacity to respond to endoderm differentiation signals both in mouse (Coronado et al. 2013) and human (Calder et al. 2013; Pauklin and Vallier 2013). In broad terms, these studies support the notion of G1 phase as a window where pluripotent stem cells can respond more effectively to differentiation cues, and specifically towards the endoderm lineage. Our findings that during the early stages of PrE differentiation cells lengthen their G1 phase as well as proliferate faster provide an explanation for how selected cells can compete with other populations occupying the same niche. In self-renewing ESCs, PrE-primed Hhex-expressing cells are slowed. Culture conditions that promote a more homogeneous expansion of an Epiblast-like state include PD03, an inhibitor of G1/S transition, suggesting that the expansion of Epiblast-like cells could also involve similar cell cycle regulation. The notion that G1 phase lengthening accompanies differentiation has been shown in other cell lineages such as neural (Lange, Huttner, and Calegari 2009; Roccio et al. 2013), pancreatic (Kim et al. 2015; Krentz et al. 2017) and intestinal (Carroll et al. 2018).

In general, we described a tendency for increased rates of proliferation accompanied by increases in G1, and hence cells exhibiting a shorter cell cycle had a longer G1. While this sort of behaviour has previously been linked to phosphorylation of Retinoblastoma protein (Rb) by ERK driving progression through the G1/S transition (ter Huurne et al. 2017), this does not fit with our observations in differentiation or with respect to PrE specification *in vivo*, where we observe a progressive increase in G1 despite the robust activation of ERK signalling. Cell cycle progression is tightly regulated by the activity of cyclin-dependent kinases (CDKs), and inhibition of cyclin E-CDK2 produces G1 lengthening and spontaneous differentiation in pluripotent stem cells (A. A. Filipczyk et al. 2007; Neganova et al. 2009; Coronado et al. 2013). In mESCs, it has been shown that MAPK/ERK pathway is a regulator of G1 length upstream of CDK/cyclin complex which phosphorylates Rb. Taking into account the heterogeneous distribution of cyclins and CDK proteins in the different cell types across development (Wianny et al. 1998), we propose that during PrE differentiation a different CDK/cyclin complex is regulating G1 phase length, and this complex is no longer a target of ERK phosphorylation in this context.

Mammalian cells share the striking quality of being synchronised in cell cycle length between members of the same generation (sisters and cousins), but not with the previous generation (mother-daughter). This was originally demonstrated in both a lymphoblast cell line and ESCs (Sandler et al. 2015; Waisman et al. 2019). Moreover, as alluded to above and suggested by previous work, this correlation might be determined based on factors that are inherited and oscillate between related cells. This means that sister cells would inherit the same amount of determinant, which would be different from the previous generation (Sandler et al. 2015). Our observations suggest that this determinant is a factor located within the MAPK pathway downstream of MEK, as culture with either the MEK inhibitor PD03 and/or the FGFR antagonist PD17 eliminates these correlations. This indicates that FGF/ERK regulation is a key nexus for this activity, and our observations suggest this activity is independent of its role in regulating differentiation. Whether this reflects a paracrine interaction of related cells with a common cytokine source or the level of receptor on the surface of these cells cannot be distinguished based on our observations.

Altogether, our data suggest that ESC cultures and their differentiation are complex models that involve an interplay of both priming-based selection and proliferation. We find that cell state changes are major drivers of culture-specific changes in proliferation that, in turn, determine the final state of a culture. Despite the increasing rates of proliferation, cells exploit G1 to integrate the signals coming from their environment and evaluate their choices. Although the time scale of the *in vivo* and *in vitro* differentiation is different, it appears that the use of increasing G1 to drive increasing levels of commitment is a fundamental facet of cell fate choice.

---

**Figure S1.**
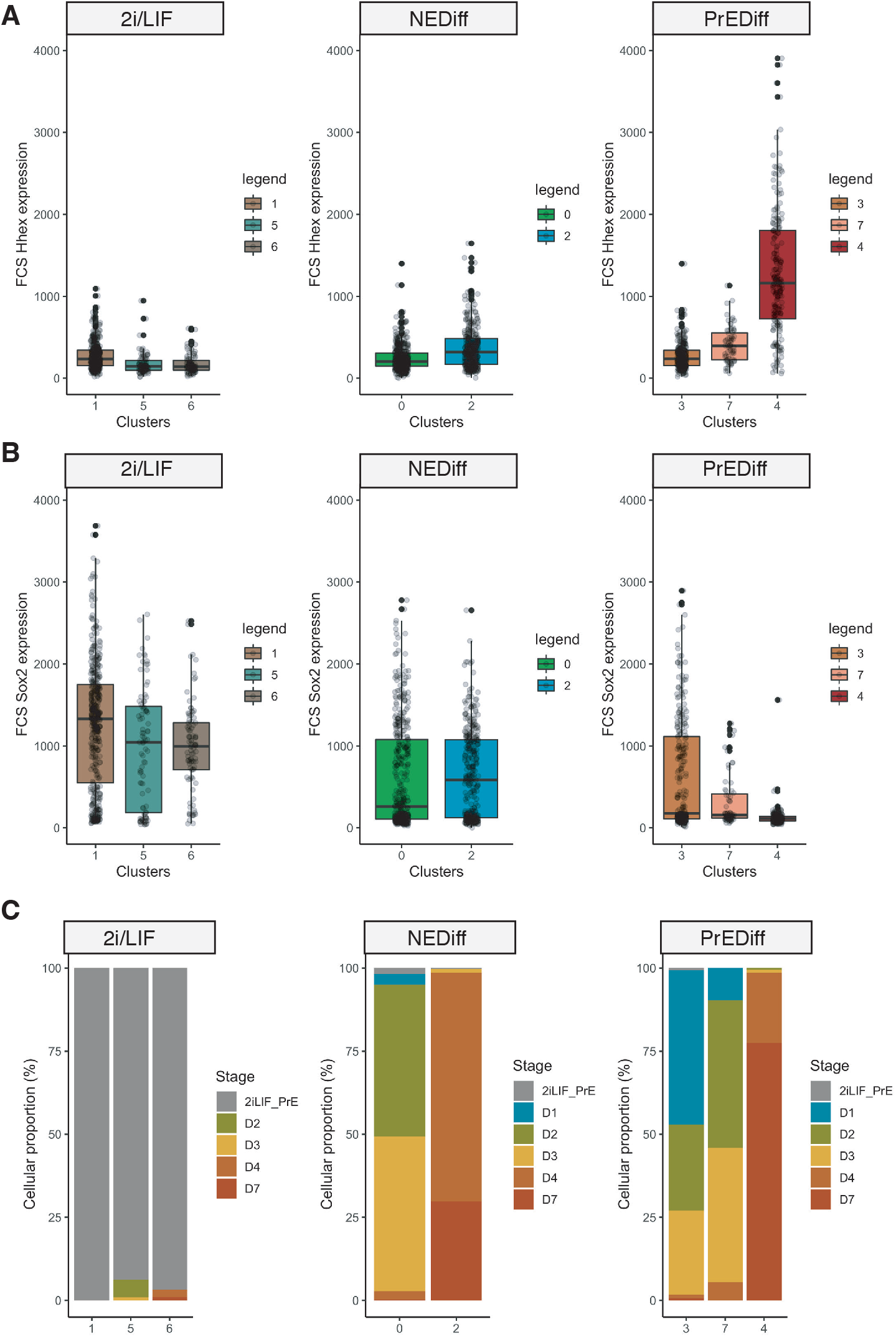
Properties of cells collected for MARS-seq. A. Fluorescence intensity on the mCherry channel recorded by the FACS at the moment of the sample collection for sequencing. B. Fluorescence intensity on the GFP channel recorded by the FACS at the moment of the sample collection for sequencing. C. Cellular proportions of cells in 2iLIF vs NEDiff vs PrE branch, showing that cells from day 2, 3 and 4 are separated between differentiated and non-differentiated.

**Figure S2.**
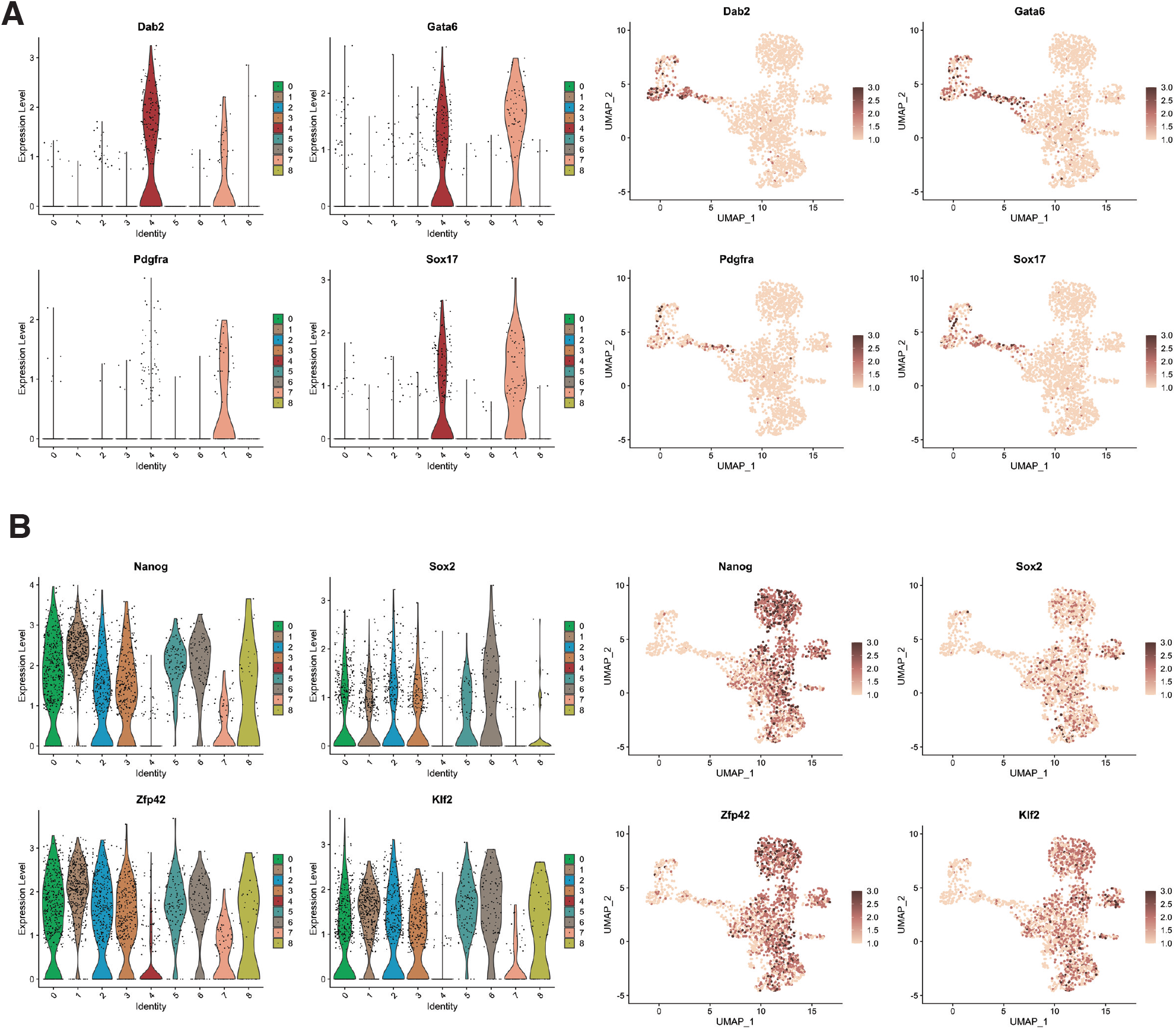
Lineage specific markers expressed in single cell RNA-seq clusters. A. Interrogation of endodermal genes (Dab2, Gata6, Pdgfra, Sox17), mostly expressed in the PrE branch of the dataset. B. Interrogation of Epiblast genes (Nanog, Sox2, Zfp42, Klf2), mostly expressed in the 2iLif clusters and the NEDiff branch of the dataset.

**Figure S3.**
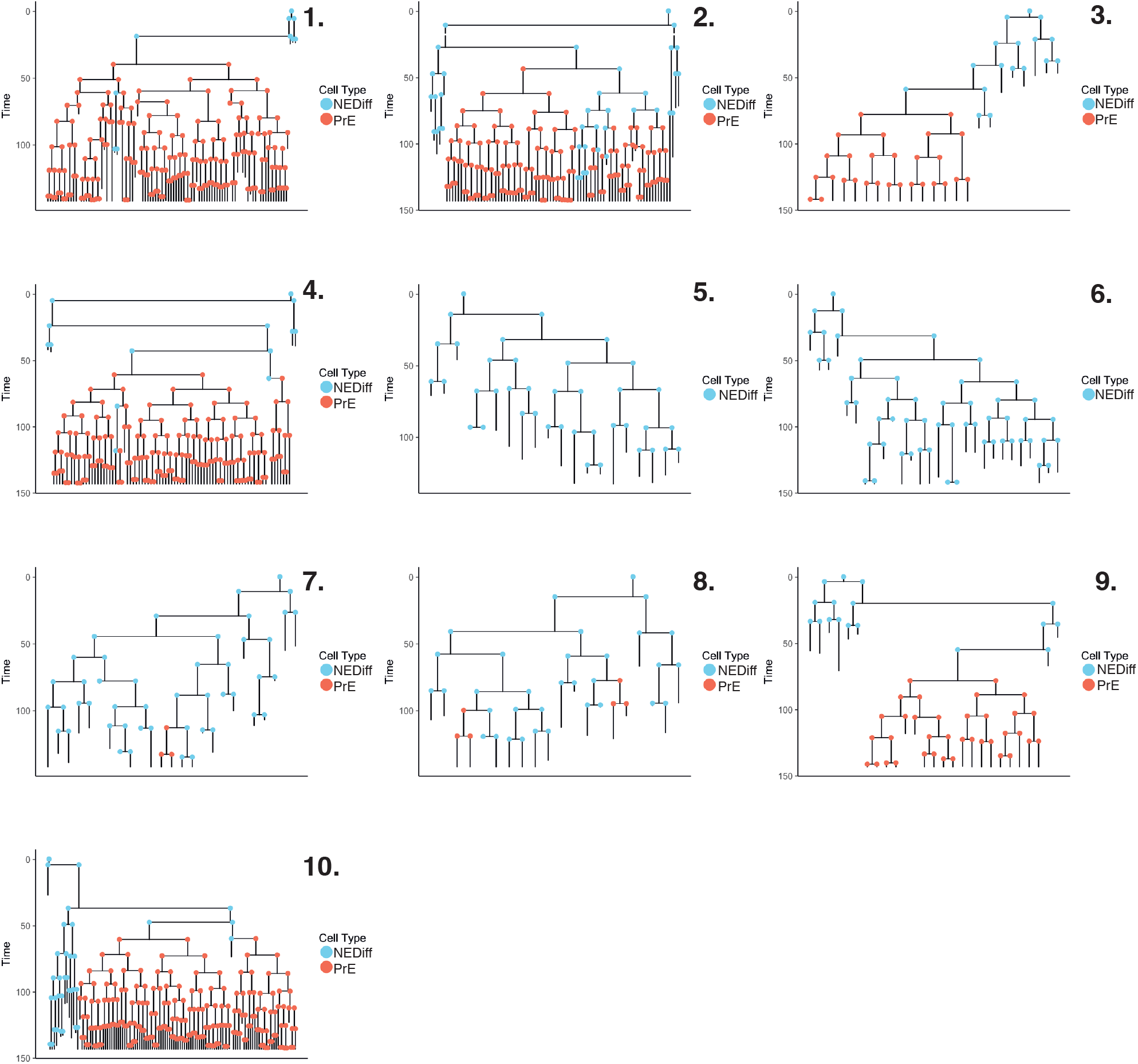
Lineage trees in PrE differentiation. Lineage trees generated in the PrE dataset. We manually tracked 1,158 cells. After constructing our lineage trees, cells that died or that had not completed a full division cycle were discarded. The final dataset consisted of 564 cells across 10 lineage trees. Cells are coloured based on clusters described in Figure 2B.

**Figure S4.**
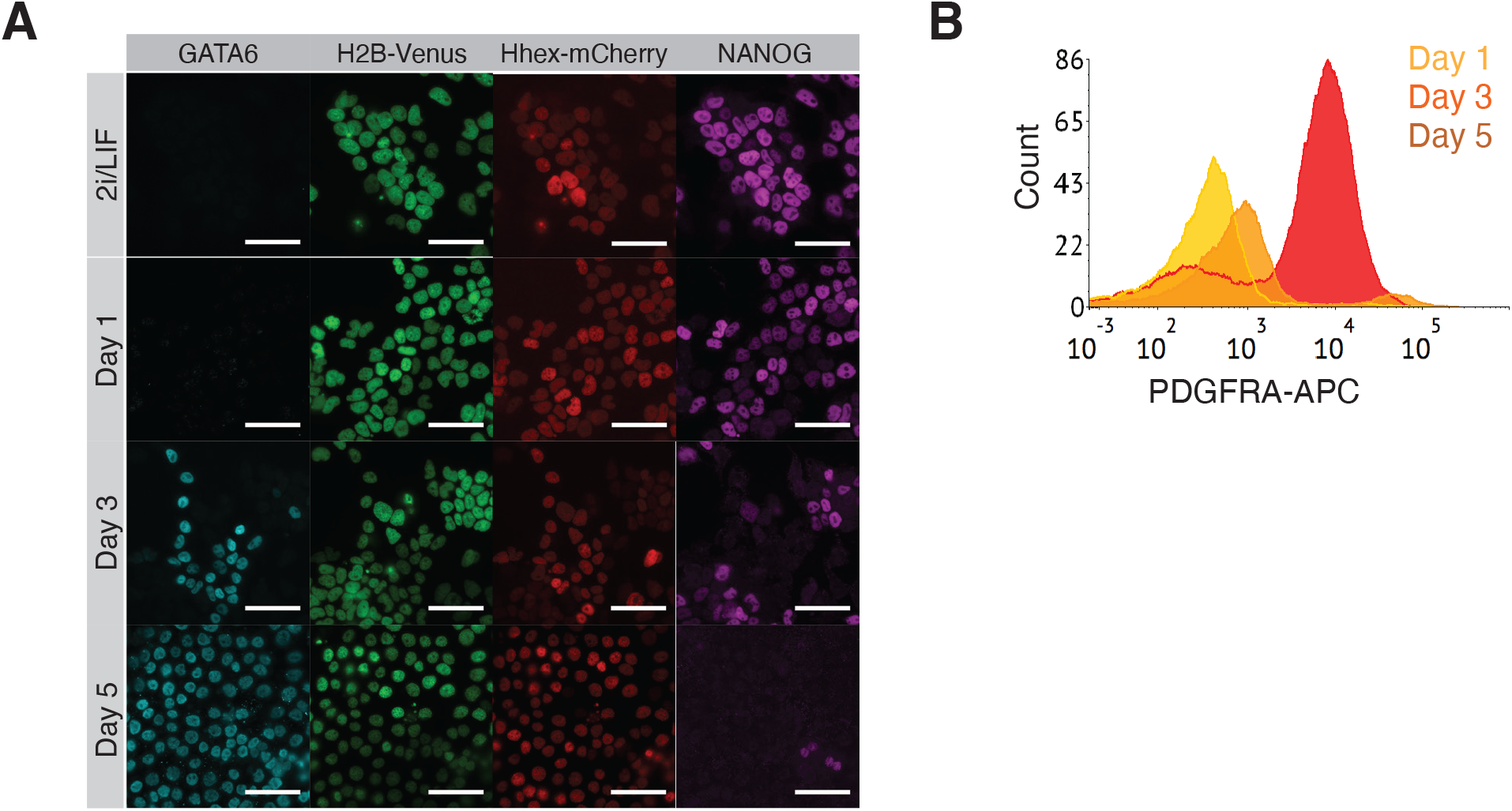
Validation of PrE differentiation. A. HFHCV mESCs lost pluripotent identity (NANOG) and acquired endodermal identity (GATA6) during PrE differentiation. Scale bar: 50 μm. B. Flow cytometry histogram of HFHCV mESCs during PrE differentiation. PDGFRA-APC staining of HFHCV mESCs at day 1,3 and 5 shows the acquisition of endodermal identity.

**Figure S5.**
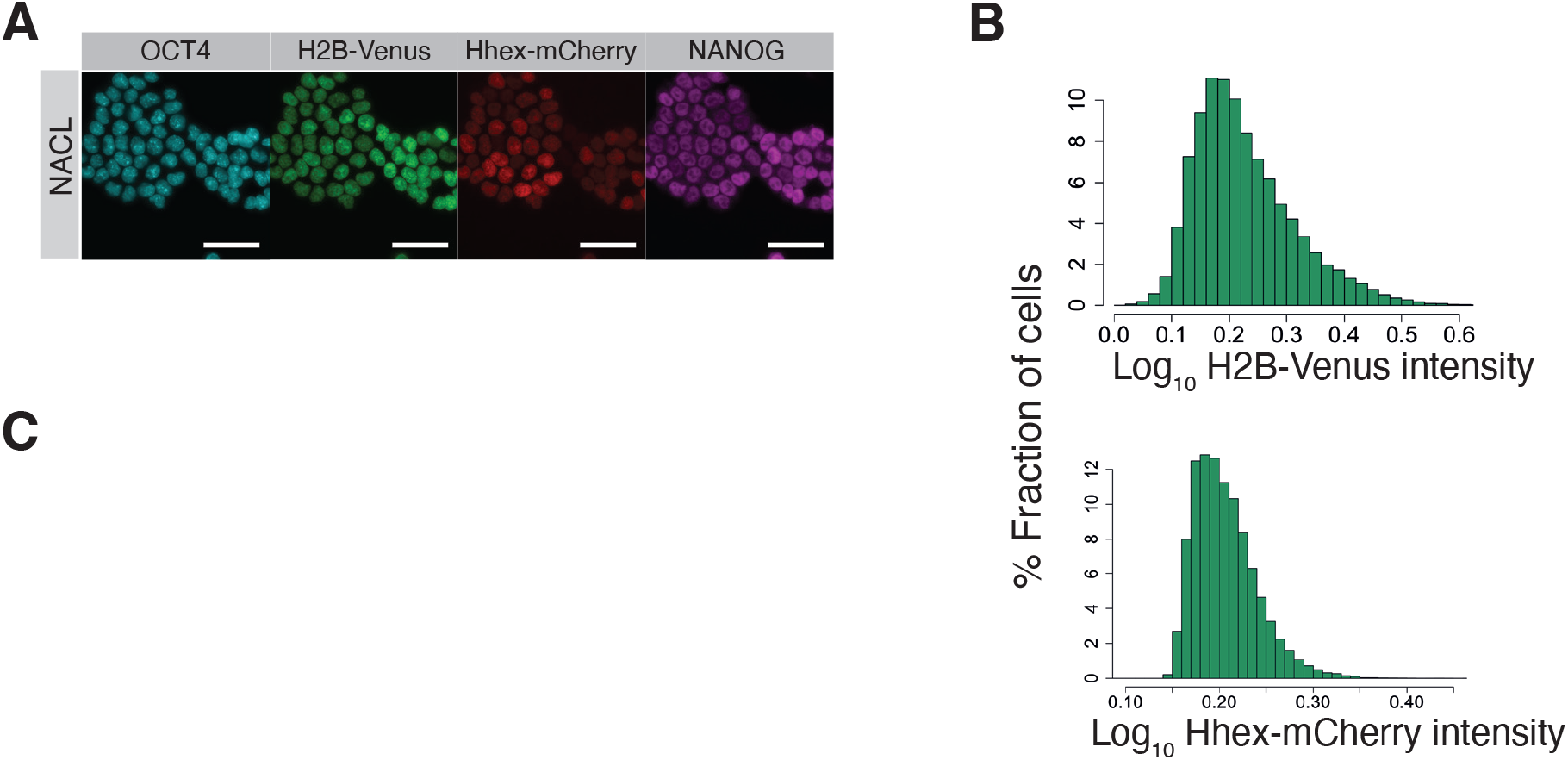
Heterogeneity in defined pluripotent stem cell culture. A. HFHCV mESCs maintained pluripotent identity after 6 days of time lapse, shown by OCT4 and NANOG immunostaining at the end of the experiment. Scale bar: 50 μm. B. Top: H2B-Venus intensity distribution for all data points collected in NACL. Not normal distribution, p-value > 0.05 in Shapiro-Wilk’s test. Bottom: Hhex-mCherry intensity distribution for all data points collected in NACL. Not normal distribution, p-value > 0.05 in Shapiro-Wilk’s test. C. Percentage of cells that transition compartments between mother and daughter cells.

**Figure S6.**
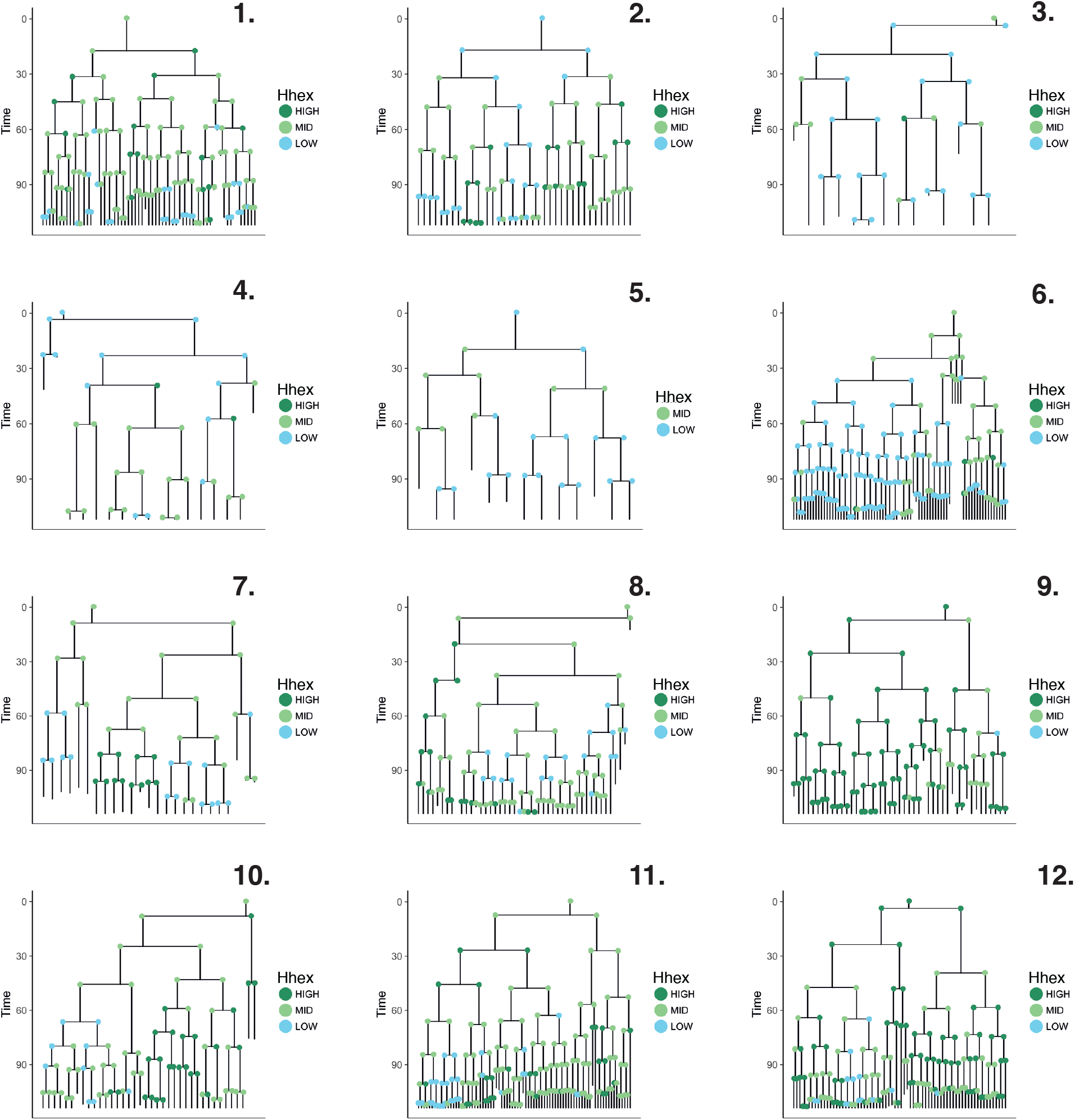
Lineage trees in pluripotent stem cell culture. Lineage trees generated in the NACL dataset. We manually tracked 1,063 cells. After constructing our lineage trees, cells that died or that had not completed a full division cycle were discarded. The final dataset consisted of 509 cells across 12 lineage trees. Cells are coloured based on *Hhex* compartments described in Figure 4C.

**Figure S7.**
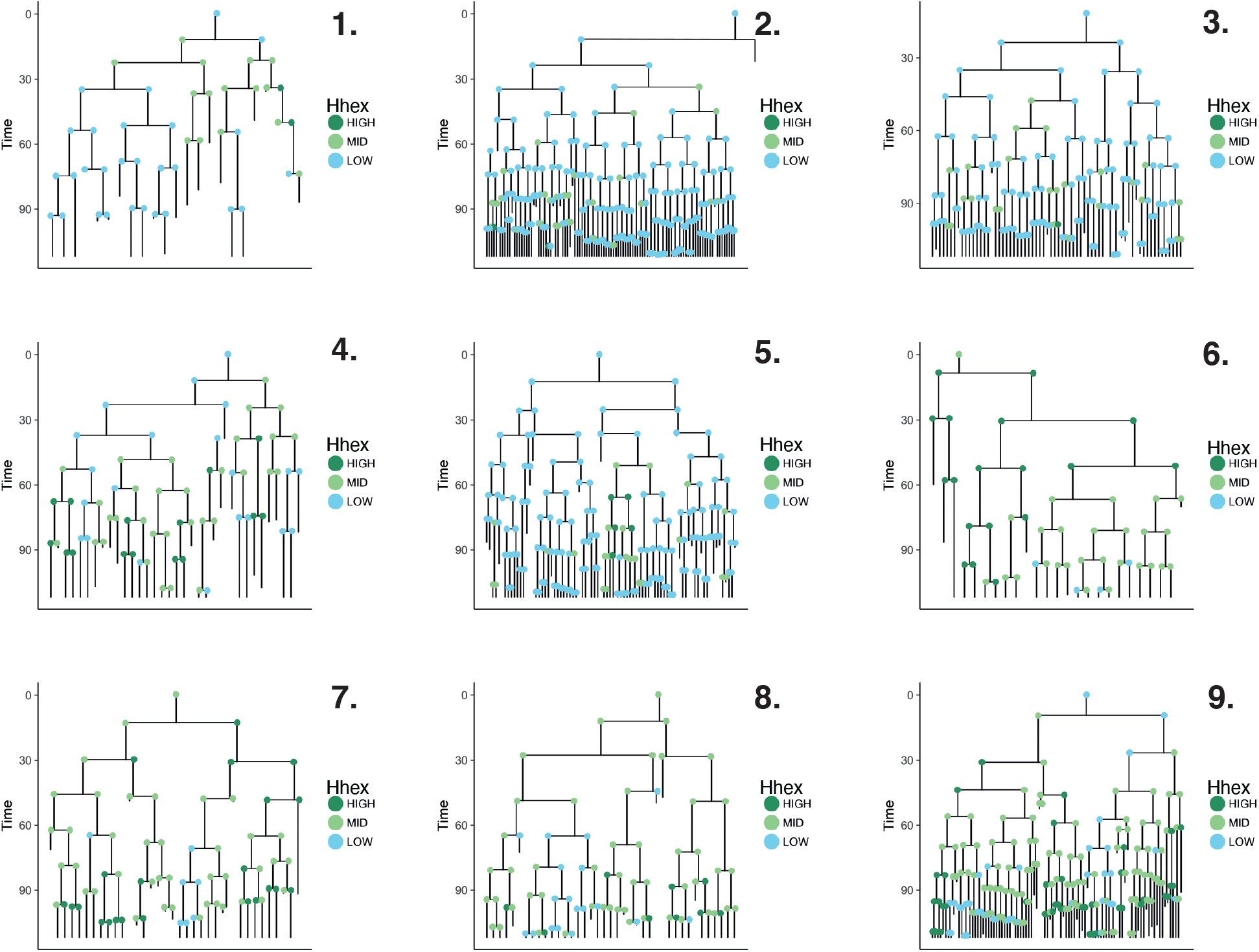
Lineage trees in pluripotent stem cell culture with PD03. Lineage trees generated in the PD03 dataset. We manually tracked 1,000 cells. After constructing our lineage trees, cells that died or that had not completed a full division cycle were discarded. The final dataset consisted of 490 cells across 9 lineage trees. Cells are coloured based on *Hhex* compartments described in Figure 4C.

**Figure S8.**
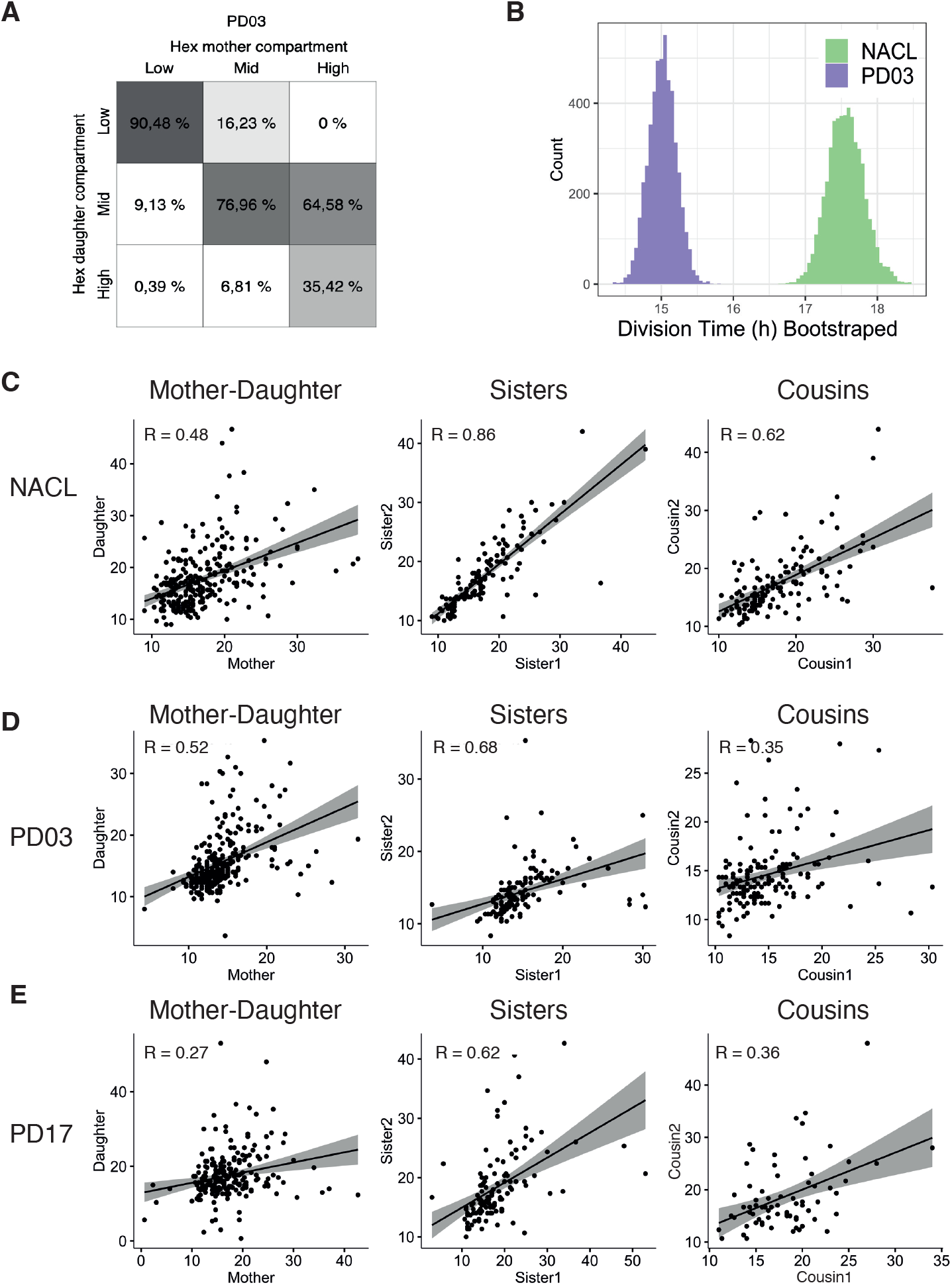
Cell cycle synchronization and its response to FGF/ERK inhibition. A. Percentage of cells that transition compartments between mother and daughter cells. B. Bootstrap analysis (100 times) of the Division Time median, showing a clear difference between the NACL and PD03 dataset. C. Spearman correlation plots for Division Times measurements in NACL. From left to right: Mother-daughter correlation, Sister pairs correlation, Cousin pairs correlation. R: Spearman Correlation Coefficient. D. Spearman correlation plots for Division Times measurements in PD03. From left to right: Mother-daughter correlation, Sister pairs correlation, Cousin pairs correlation. R: Spearman Correlation Coefficient. E. Spearman correlation plots for Division Times measurements in PD17. From left to right: Mother-daughter correlation, Sister pairs correlation, Cousin pairs correlation. R: Spearman Correlation Coefficient.

**Figure S9.**
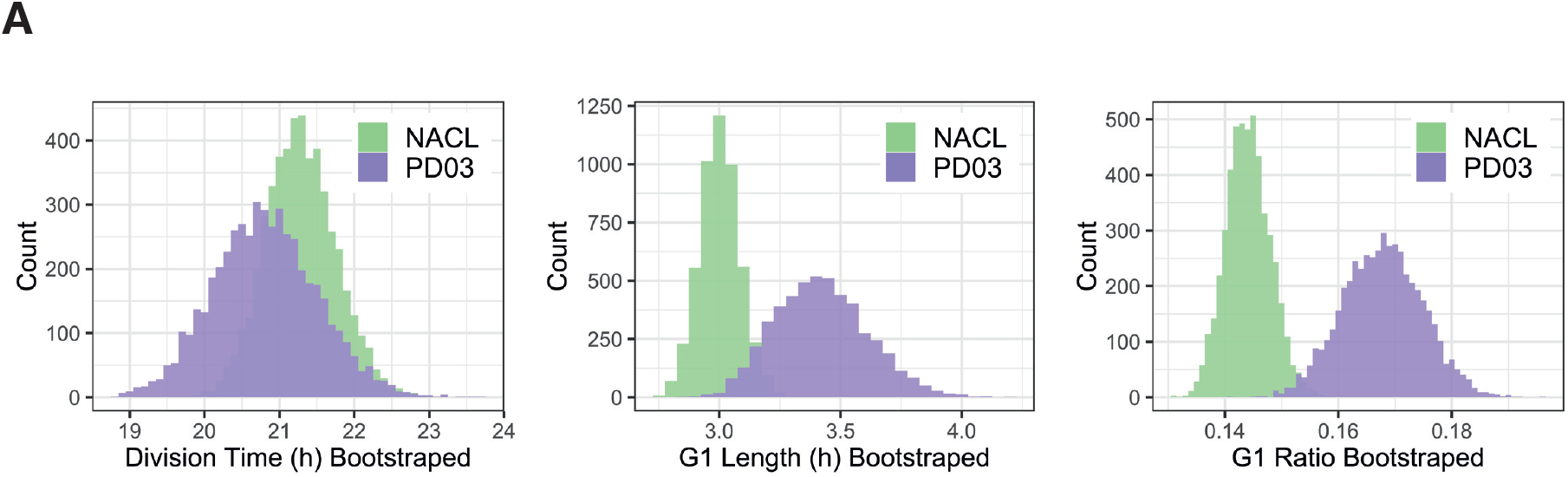
Bootstrap analysis of the FUCCI dataset. A. Bootstrap analysis (100 times) of Division Time, G1 Length and G1 Ratio measures in the FUCCI cell line, showing a difference between the NACL and PD03 dataset.

**Table S1.** Summary of the cell numbers in the different scRNA-seq clusters analysed. Clusters are annotated as NEDiff, PrE Parental and PrE Diff according to Fig 1B.

|  |  | 2i/LIF | Day 1 | Day 2 | Day 3 | Day 4 | Day 7 |
| --- | --- | --- | --- | --- | --- | --- | --- |
| 2i/LIF | Cluster 1 | 415 | 0 | 0 | 0 | 0 | 0 |
|  | Cluster 5 | 105 | 0 | 6 | 1 | 0 | 0 |
|  | Cluster 6 | 91 | 0 | 0 | 0 | 2 | 1 |
| NEDiff | Cluster 0 | 8 | 14 | 201 | 205 | 11 | 1 |
|  | Cluster 2 | 1 | 0 | 0 | 4 | 238 | 103 |
| PrE Parental | Cluster 3 | 2 | 139 | 78 | 76 | 3 | 2 |
| PrE Diff | Cluster 7 | 0 | 7 | 32 | 29 | 4 | 0 |
|  | Cluster 4 | 0 | 0 | 1 | 2 | 44 | 162 |
|  | Total | 622 | 160 | 318 | 317 | 302 | 269 |

**Table S2.** Comparison of survival and proliferation during time lapse of PrE differentiation. Survival rate is calculated as a ratio between cells that survived between total cells. Death rate is the ratio between cells that died and total number of cells. As total cells, only cells with complete cell cycle information are considered. Division Time (hours) is shown as median ± standard deviation.

|  | PrEDiff |  |  |  | NEDiff |  |  |  |  |
| --- | --- | --- | --- | --- | --- | --- | --- | --- | --- |
|  | Day 2 | Day 3 | Day 4 | Day 5 | Day 1 | Day 2 | Day 3 | Day 4 | Day 5 |
| Cells that survive | 5 | 32 | 102 | 228 | 54 | 41 | 38 | 42 | 21 |
| Cells that die | 1 | 4 | 7 | 24 | 20 | 42 | 16 | 40 | 57 |
| Cells total | 6 | 36 | 109 | 252 | 74 | 83 | 54 | 82 | 78 |
| Survival rate | 0.83 | 0.89 | 0.94 | 0.90 | 0.73 | 0.49 | 0.70 | 0.51 | 0.27 |
| Death rate | 0.17 | 0.11 | 0.06 | 0.10 | 0.27 | 0.51 | 0.30 | 0.49 | 0.73 |
| Division Time (h) | 13.33 ± 4 | 11.00 ± 3 | 13.00 ± 4 | 17.33 ± 4 | 16.33 ± 4 | 18.00 ± 7 | 20.00 ± 6 | 17.00 ± 7 | 22.00 ± 7 |

**Table S3.** Analysis of PrE parental cells. Division Time (hours) is shown as median ± standard deviation. Hhex-mCherry fluorescence (absolute units) is shown as median ± standard deviation.

|  | PrEFate | NotPrEFate |
| --- | --- | --- |
| Cells | 20 | 151 |
| Division Time (h) | 17.67 ± 6 | 17.33 ± 6 |
| Hhex-mCherry (a.u.) | 0.18 ± 0.03 | 0.15 ± 0.02 |

**Table S4.** Comparison of survival and proliferation between Hhex compartments. Survival rate is produced as a ratio between cells that survived between total cells. Death rate is the ratio between cells that died and total number of cells. As total cells, only cells with complete cell cycle information are considered. Division Time (hours) is shown as median ± standard deviation.

|  | NACL |  |  | PD03 |  |  |
| --- | --- | --- | --- | --- | --- | --- |
|  | HIGH | MID | LOW | HIGH | MID | LOW |
| Cells that survive | 127 | 254 | 128 | 48 | 191 | 252 |
| Cells that die | 12 | 35 | 34 | 14 | 55 | 55 |
| Cells total | 139 | 289 | 162 | 62 | 246 | 307 |
| Survival rate | 0.91 | 0.88 | 0.79 | 0.77 | 0.78 | 0.82 |
| Death rate | 0.09 | 0.12 | 0.21 | 0.23 | 0.22 | 0.18 |
| Division Time (h) | 17.33 ± 6 | 16.33 ± 6 | 14.67 ± 5 | 15.83 ± 5 | 14.00 ± 4 | 13.67 ± 4 |

**Table S5.** Summary of dataset collected in the FUCCI cell line. Division Time (hours) is shown as median ± standard deviation. G1 Length (hours) is shown as median ± standard deviation. G1 Ratio is produced as the ratio between the G1Time and the total Division Time, and it is shown as median ± standard deviation.

|  | NACL | PD03 | 2i/LIF | PrE |  |  |
| --- | --- | --- | --- | --- | --- | --- |
|  |  |  |  | Day 2 | Day 3 | Day 4 |
| Division Time (h) | 20.4 ± 5 | 19.5 ± 7 | 15.0 ± 5 | 16.33 ± 1 | 15.0 ± 5 | 18.0 ± 3 |
| G1 Length (h) | 2.7 ± 2.5 | 3.1 ± 1.9 | 3.0 ± 0.9 | 2.7 ± 1.2 | 4.3 ± 1.5 | 7.3 ± 2.9 |
| G1 Ratio | 0.14 ± 0.1 | 0.15 ± 0.1 | 0.19 ± 0.1 | 0.16 ± 0.1 | 0.24 ± 0.1 | 0.40 ± 0.1 |
| Cells total | 126 | 104 | 110 | 12 | 29 | 39 |

## Supplementary Movies Legends

**Supplementary Movie 1. Example of a tracked HFHCV colony during PrE differentiation.**

Cyan is *H2B**-**Venus*, magenta is *Hhex-mCherry*. Yellow squares show the cell tracking. Scale bar is 300 μm.

**Supplementary Movie 2. Example of a tracked HFHCV colony in NACL.**

*H2B-Venus* is shown in cyan, magenta is *Hhex-mCherry*. Yellow squares show the cell tracking. Scale bar is 200 μm.

**Supplementary Movie 3. Example of an imaged FUCCI colony in NACL.**

*mCherry-Cdtl* is shown in cyan, and *H2B-miRF670* is shown in magenta. Scale bar is 20 μm.

**Supplementary Movie 4. Example of an imaged FUCCI colony during PrE differentiation.**

*mCherry-Cdt1* is shown in cyan, and *H2B-miRF670* is shown in magenta. Scale bar is 300 μm.

## Methods

### Mouse ESC Culture and cell lines

Mouse ESC lines were cultured in standard conditions in NACL as previously described in Anderson et al., 2017. For NACL+PD03 experiments, PD 0325901 (PD03) was added at the final concentration of 1μM. For PrE experiments, cells were cultured previously in 2i/LIF containing CHI 99021 (Chiron), PD03 and LIF, and then in RACL as described in Anderson et al., 2017.

The cell line HFHCV was previously reported in (Illingworth et al. 2016). It contains Hhex-3xFLAG-IRES-H2b-mCherry (HFHCVz), and a pCAG-H2b-Venus vector (HFHCV). Using this cell line *Hhex* transcription can be visualized by translational amplification of *Hhex*, which drives the expression of the monomeric fluorescent protein mCherry (Canham et al. 2010; S. M. Morgani et al. 2013; Illingworth et al. 2016).

The FUCCI-H2B-miRF670 reporter construct was constructed as follows. The H2B and miRFP670 sequences were fused by overlap extension PCR. The miRFP670 sequence was cloned from pY42-pmiRFP670AAA-NLS-Myc, a kind gift from YH Kim. The resulting fragment H2B-miRFP670 was inserted into the PCR-Blunt II-TOPO backbone by TOPO cloning. The H2B-miRFP670 fragment was digested with the *Cpol* (*Rsrll*) and *Klfl* enzymes and inserted upstream of the Hygromycin resistance gene cassette of the ES-FUCCI plasmid (Addgene #62451). RAF-ER^T2^ (Hamilton and Brickman 2014) cells were electroporated with *BgII*-linearised FUCCI-H2B-miRFP670 DNA (25μg), and stable transfectants were selected for with hygromycin (125μg/ml).

### Time lapse

Mouse ESC lines were cultured in NACL or 2i/LIF medium on Laminin 511(BioLamina) -coated 8-well slides (Ibidi) and imaged at 20 minutes intervals for 6 days in mCherry and Venus fluorescent light channels, in 5%CO_2_ and 20%O_2_ at 37°C under a Deltavision Widefield Screening microscope. ESCs were seeded at 300 cells/cm^2^ 48h before the beginning of the time lapse for NACL and PD03 experiments, or at 30.000 cells/cm^2^ 24h before for PrE differentiation. For PD03 experiments, cells were cultured in NACL for at least 2 passages and PD03 was added just before starting the time lapse. For PrE experiments, cells were cultured in 2i/LIF for 2 passages and then changed to RPMI minimal medium just before starting the experiment. Activin A, Chiron and LIF (RACL) were added 24h after the time lapse started. In all experiments, the media was changed every day of the time lapse. To test that the laminin coating was not affecting the behaviour of the HFHCV cell line, we sorted Hhex-high and low populations seeded in both gelatine and laminin and analysed them 24 and 48h later to test that their re-equilibration rates are the same.

### Flow cytometry

Cells were collected by trypsinisation and stained against a marker of undifferentiated mESCs, Pecam-1 (BD Biosciences, APC-conjugated, 551262; 1:200), or a marker for PrE differentiated cells, PDGFRA (BD Biosciences, APC-conjugated, 562777; 1:200), and DAPI (Molecular Probes, D1306, 1 μg/ml) to exclude dead cells. mESCs were stained for 15 min at 4°C before being washed and resuspended in FACS buffer (10%FCS in PBS) with DAPI. Flow cytometry analysis was carried out using a BD LSR Fortessa, and flow cytometry sorting was carried out in a BD FACS Aria III. Data analysis was carried out using FCS Express 6 Flow software (De Novo Software) by gating on forward and side scatter to identify a cell population and eliminate debris, then gating DAPI negative, viable cells before assessing the levels of GFP, mCherry or APC.

### mESCs immunostaining

Mouse ESCs were cultured in 8-wells slides (Ibidi). ESC immunostaining was carried out as previously described in Canham et al., 2010. The following antibodies were used (all at 1:200): anti-NANOG (eBioscience, 14-5761), anti-OCT4 (Santa Cruz, sc-5279), anti-GATA6 (Cell Signalling Technologies, 5851). Secondary antibodies used are from the Alexa fluor series (Molecular Probes, ThermoFisher). mESCs were imaged using a Deltavision Widefield Screening microscope.

### Cell tracking

We performed manual cell tracking using Imaris v9.5 (Bitplane). Nuclei were segmented using the H2B marker, and we measured the Hhex-mCherry fluorescence intensity of a circular area of 50 μm diameter inside the segmented nuclei. For each area measured, we took the median fluorescence intensity as the measure for that given data point. Intensity measurements were linked to its time point and lineage, allowing us to infer the division time for each cell that was tracked, as well as the expression level of *Hhex* in each time point. Only cells with completed cell cycle information were used for cell cycle and compartment analysis.

### Data Analysis

Data mining was performed with Matlab. A script was generated to take the separated output data frames from Imaris and convert into a single file containing all the dataset information, by cell. This file also contains the lineage tree information, mean fluorescence intensities and division times. Alive and dead cells, as well as cells without complete cell cycle information, are located into groups for further analysis. Another file organised by time frame is generated in order to perform time dynamics analysis.

Statistical analysis and plotting were performed in R. *Hhex* compartments were determined by dividing the Hhex-mCherry intensity distribution into quartiles. All cells above the first quartile (0.1648) are considered High, and all cells below the third quartile (0.1376) are Low. The same quartiles are used for the PD03 dataset (Table S4). The PrE dataset was divided into PrE cluster and NEDiff cluster using *k*-means clustering.

Plots were created with R ggplot2 package, and GraphPad Prism.

### Mathematical modelling in Figure 3

The model starts with *N* cells of cell type 1 (called generation 1). For each subsequent generation, all cells divide into two new cells. Each new cell undergoes a transition with probability *t_12_* to become cell type 2. In each generation, all cells of cell type 1 survive with probability *s_1_* and all cells of cell type 2 survive with probability *s_2_*. All cells inherit their parent cell’s cell type and cannot transition back to cell type 1. This model continues for 8 generations corresponding to almost 6 days (generation time = 17 hours). With no cell death rate, this model results in exponential growth.

### scRNA-seq analysis

Single cell libraries were sequenced using Illumina NextSeq 500. Pre-processing steps and quality control was done as described previously in (Jaitin et al. 2014; Rothová et al. 2022). In short, the reads were aligned using HISAT (version 0.1.6) and mapped to mouse mm9 genome. Further downstream analysis was done using scanpy (version 1.4.6). After filtering 2,028 cells and 17,769 genes were normalized and log-transformed. We identified 2,000 highly variable genes using seurat flavor. Genes were further scaled to mean-variance followed by PCA and UMAP dimension reduction. No cell cycle regression and batch correction were necessary. Finally, we used unsupervised Louvain clustering with resolution set to 0.8 and which identified 9 overall clusters (04_analysis.Rmd). Cell cycle was estimated using Seurat’s (4.0.1) *CellCycleScoring* function.

### Cluster Alignment Tool (CAT)

To estimate similarity between clusters we used in-house CAT. First, the tool normalized the datasets using non-zero median which removes the influence of outlier genes. Both datasets are subset for unique and common genes. Next, Euclidean distance is calculated between each pair of clusters by randomly sampling cells with replacement to cluster size. This step is repeated 1,000 times generating two distributions. If the distance is significant (sigma = 1.6 representing p-val 0.05) we define these clusters transcriptionally similar.

## Code availability

We deposited the original MATLAB scripts behind the data mining and mathematical models on Github: https://github.com/SilasBoyeNissen/Link-between-cell-cycle-length-regulation-and-endoderm-priming-in-vitro

All scRNA-seq analysis was also uploaded to Github at: http://github.com/brickmanlab/perera_silas_et_al_2022/.

## Data Availability

The sc-RNAseq data used in this study have been deposited in the Gene Expression Omnibus, the accession number will be available shortly and all data will be made publicly accessible. Previously published Nowotschin et al., 2019 data that were used here are available under accession code GSE123046.

## Materials Availability

All materials will be available upon reasonable request to the corresponding author.

## Acknowledgements

We thank James Briscoe for critical discussion, Jose Alejandro Herrera Romero for bioinformatics advice, YH Kim for the RFP670 plasmid, and the Brickman laboratory members for critical discussion and reading of this manuscript. We thank Helen Neil, Magali Michaut and the reNEW Genomics Platform for technical expertise, support, and the use of instruments. We thank Jutta Bulkescher and the reNEW Imaging Platform for training, technical expertise, support, and the use of microscopes. We thank Gelo dela Cruz, Paul van Dieken and the reNEW Flow Cytometry Platform for technical expertise, support, and the use of instruments. This work was supported by grants from the Lundbeck Foundation (R198-2015-412); Independent Research Fund Denmark (DFF-8020-00100B); the Danish National Research Foundation (DNRF116); M.P was supported by a PhD studentship from the Lundbeck Foundation (R286-2018-1534); R.S.M. was supported by a fellowship from the Lundbeck Foundation (R303-2018-2939); The Novo Nordisk Foundation Center for Stem Cell Medicine (reNEW) is supported by a Novo Nordisk Foundation grant, number NNF21CC0073729; the Novo Nordisk Foundation Center for Stem Cell Biology, was supported by a Novo Nordisk Foundation grant, number NNF17CC0027852.

## References

Abranches, Elsa, Ana M.V. Guedes, Martin Moravec, Hedia Maamar, Petr Svoboda, Arjun Raj, and Domingos Henrique. 2014. ‘Stochastic NANOG Fluctuations Allow Mouse Embryonic Stem Cells to Explore Pluripotency’. Development (Cambridge) 141 (14): 2770–79. https://doi.org/10.1242/dev.108910.

Anderson, Kathryn G.V., William B. Hamilton, Fabian V. Roske, Ajuna Azad, Teresa E. Knudsen, Maurice A. Canham, Lesley M. Forrester, and Joshua M. Brickman. 2017. ‘Insulin Fine-Tunes Self-Renewal Pathways Governing Naive Pluripotency and Extra-Embryonic Endoderm’. Nature Cell Biology 19 (10): 1164–77. https://doi.org/10.1038/ncb3617.

Beddington, R.S., and E.J. Robertson. 1989. ‘An Assessment of the Developmental Potential of Embryonic Stem Cells in the Midgestation Mouse Embryo’. Development 105 (4): 733–37. https://doi.org/10.1242/dev.105.4.733.

Calder, Ashley, Ivana Roth-Albin, Sonam Bhatia, Carlos Pilquil, Jong Hee Lee, Mick Bhatia, Marilyne Levadoux-Martin, et al. 2013. ‘Lengthened G1 Phase Indicates Differentiation Status in Human Embryonic Stem Cells’. Stem Cells and Development 22 (2): 279–95. https://doi.org/10.1089/scd.2012.0168.

Canham, Maurice A., Alexei A. Sharov, Minoru S.H. Ko, and Joshua M. Brickman. 2010. ‘Functional Heterogeneity of Embryonic Stem Cells Revealed through Translational Amplification of an Early Endodermal Transcript’. PLoS Biology 8 (5). https://doi.org/10.1371/journal.pbio.1000379.

Cannon, D., A. M. Corrigan, A. Miermont, P. McDonel, and J. R. Chubb. 2015. ‘Multiple Cell and Population-Level Interactions with Mouse Embryonic Stem Cell Heterogeneity’. Development 142 (16): 2840–49. https://doi.org/10.1242/dev.120741.

Carroll, Thomas D., Ian P. Newton, Yu Chen, J. Julian Blow, and Inke Näthke. 2018. ‘Lgr5+ Intestinal Stem Cells Reside in an Unlicensed G1 Phase’. Journal of Cell Biology 217 (5): 1667–85. https://doi.org/10.1083/jcb.201708023.

Chazaud, Claire, Yojiro Yamanaka, Tony Pawson, and Janet Rossant. 2006. ‘Early Lineage Segregation between Epiblast and Primitive Endoderm in Mouse Blastocysts through the Grb2-MAPK Pathway’. Developmental Cell 10 (5): 615–24. https://doi.org/10.1016/j.devcel.2006.02.020.

Coronado, Diana, Murielle Godet, Pierre-Yves Bourillot, Yann Tapponnier, Agnieszka Bernat, Maxime Petit, Marielle Afanassieff, et al. 2013. ‘A Short G1 Phase Is an Intrinsic Determinant of Naïve Embryonic Stem Cell Pluripotency’. Stem Cell Research 10 (1): 118–31. https://doi.org/10.1016/j.scr.2012.10.004.

Evans, M.J., and M. H. Kaufman. 1981. ‘Establishment in Culture of Pluripotential Cells from Mouse Embryos’. Nature 292 (July): 154–56.

Filipczyk, Adam A., Andrew L. Laslett, Christine Mummery, and Martin F. Pera. 2007. ‘Differentiation Is Coupled to Changes in the Cell Cycle Regulatory Apparatus of Human Embryonic Stem Cells’. Stem Cell Research 1 (1): 45–60. https://doi.org/10.1016/j.scr.2007.09.002.

Filipczyk, Adam, Carsten Marr, Simon Hastreiter, Justin Feigelman, Michael Schwarzfischer, Philipp S. Hoppe, Dirk Loeffler, et al. 2015. ‘Network Plasticity of Pluripotency Transcription Factors in Embryonic Stem Cells’. Nature Cell Biology 17 (10): 1235–46. https://doi.org/10.1038/ncb3237.

Hamilton, William B., and Joshua M. Brickman. 2014. ‘Erk Signaling Suppresses Embryonic Stem Cell Self-Renewal to Specify Endoderm’. Cell Reports 9 (6): 2056–70. https://doi.org/10.1016/j.celrep.2014.11.032.

Hamilton, William B., Yaron Mosesson, Rita S. Monteiro, Kristina B. Emdal, Teresa E. Knudsen, Chiara Francavilla, Naama Baarkai, Jesper V. Olsen, and Joshua M. Brickman. 2019. ‘Dynamic Lineage Priming Is Driven via Direct Enhancer Regulation by ERK’. Nature 575: 355–60. https://doi.org/10.1038/s41586-019-1732-z.

Hastreiter, Simon, Stavroula Skylaki, Dirk Loeffler, Andreas Reimann, Oliver Hilsenbeck, Philipp S. Hoppe, Daniel L. Coutu, et al. 2018. ‘Inductive and Selective Effects of GSK3 and MEK Inhibition on Nanog Heterogeneity in Embryonic Stem Cells’. Stem Cell Reports 11 (1): 58–69. https://doi.org/10.1016/j.stemcr.2018.04.019.

Huurne, Menno ter, James Chappell, Stephen Dalton, and Hendrik G. Stunnenberg. 2017. ‘Distinct Cell-Cycle Control in Two Different States of Mouse Pluripotency’. Cell Stem Cell 21 (4): 449–455.e4. https://doi.org/10.1016/j.stem.2017.09.004.

Illingworth, Robert S, Jurriaan J Hölzenspies, Fabian V Roske, Wendy A Bickmore, and Joshua M Brickman. 2016. ‘Polycomb Enables Primitive Endoderm Lineage Priming in Embryonic Stem Cells’. ELife 5: 1–28. https://doi.org/10.7554/elife.14926.

Jaitin, Diego Adhemar, Ephraim Kenigsberg, Hadas Keren-Shaul, Naama Elefant, Franziska Paul, Irina Zaretsky, Alexander Mildner, et al. 2014. ‘Massively Parallel Single-Cell RNA-Seq for Marker-Free Decomposition of Tissues into Cell Types’. Science 343 (6172): 776–79. https://doi.org/10.1126/science.1247651.

Kim, Yung Hae, Hjalte List Larsen, Pau Rué, Laurence A. Lemaire, Jorge Ferrer, and Anne Grapin-Botton. 2015. ‘Cell Cycle–Dependent Differentiation Dynamics Balances Growth and Endocrine Differentiation in the Pancreas’. PLoS Biology 13 (3): e1002111. https://doi.org/10.1371/journal.pbio.1002111.

Krentz, Nicole A. J., Dennis van Hoof, Zhongmei Li, Akie Watanabe, Mei Tang, Cuilan Nian, Michael S. German, and Francis C. Lynn. 2017. ‘Phosphorylation of NEUROG3 Links Endocrine Differentiation to the Cell Cycle in Pancreatic Progenitors’. Developmental Cell 41 (2): 129–142.e6. https://doi.org/10.1016/j.devcel.2017.02.006.

Kunath, Tilo, Marc K. Saba-El-Leil, Marwa Almousailleakh, Jason Wray, Sylvain Meloche, and Austin Smith. 2007. ‘FGF Stimulation of the Erk1/2 Signalling Cascade Triggers Transition of Pluripotent Embryonic Stem Cells from Self-Renewal to Lineage Commitment’. Development 134 (16): 2895–2902. https://doi.org/10.1242/dev.02880.

Lallemand, Y., and P. Brulet. 1990. ‘An in Situ Assessment of the Routes and Extents of Colonisation of the Mouse Embryo by Embryonic Stem Cells and Their Descendants’. Development 110 (4): 1241–48. https://doi.org/10.1242/dev.110.4.1241.

Lange, Christian, Wieland B. Huttner, and Federico Calegari. 2009. ‘Cdk4/CyclinD1 Overexpression in Neural Stem Cells Shortens G1, Delays Neurogenesis, and Promotes the Generation and Expansion of Basal Progenitors’. Cell Stem Cell 5 (3): 320–31. https://doi.org/10.1016/j.stem.2009.05.026.

Martello, Graziano, and Austin Smith. 2014. ‘The Nature of Embryonic Stem Cells’. Annual Review of Cell and Developmental Biology 30 (1): 647–75. https://doi.org/10.1146/annurev-cellbio-100913-013116.

Martin, G R. 1981. ‘Isolation of a Pluripotent Cell Line from Early Mouse Embryos Cultured in Medium Conditioned by Teratocarcinoma Stem Cells.’ Proceedings of the National Academy of Sciences of the United States of America 78 (12): 7634–38.

Morgani, Sophie M., Maurice A. Canham, Jennifer Nichols, Alexei A. Sharov, Rosa Portero Migueles, Minoru S.H. Ko, and Joshua M. Brickman. 2013. ‘Totipotent Embryonic Stem Cells Arise in Ground-State Culture Conditions’. Cell Reports 3 (6): 1945–57. https://doi.org/10.1016/j.celrep.2013.04.034.

Morgani, Sophie, Jennifer Nichols, and Anna-Katerina Hadjantonakis. 2017. ‘The Many Faces of Pluripotency: In Vitro Adaptations of a Continuum of in Vivo States’. BMC Developmental Biology 17 (1): 7. https://doi.org/10.1186/s12861-017-0150-4.

Mummery, C. L., C. E. van den Brink, and S. W. de Laat. 1987. ‘Commitment to Differentiation Induced by Retinoic Acid in P19 Embryonal Carcinoma Cells Is Cell Cycle Dependent’. Developmental Biology 121 (1): 10–19. https://doi.org/10.1016/0012-1606(87)90133-3.

Neganova, I., X. Zhang, S. Atkinson, and M. Lako. 2009. ‘Expression and Functional Analysis of G1 to S Regulatory Components Reveals an Important Role for CDK2 in Cell Cycle Regulation in Human Embryonic Stem Cells’. Oncogene 28 (1): 20–30. https://doi.org/10.1038/onc.2008.358.

Nichols, Jennifer, Jose Silva, Mila Roode, and Austin Smith. 2009. ‘Suppression of Erk Signalling Promotes Ground State Pluripotency in the Mouse Embryo’. Development 136 (19): 3215–22. https://doi.org/10.1242/dev.038893.

Nowotschin, Sonja, Manu Setty, Ying-Yi Kuo, Vincent Liu, Vidur Garg, Roshan Sharma, Claire S. Simon, et al. 2019. ‘The Emergent Landscape of the Mouse Gut Endoderm at Single-Cell Resolution’. Nature 569 (7756): 361–67. https://doi.org/10.1038/s41586-019-1127-1.

Pauklin, Siim, and Ludovic Vallier. 2013. ‘The Cell-Cycle State of Stem Cells Determines Cell Fate Propensity’. Cell 155 (1): 135–47. https://doi.org/10.1016/j.cell.2013.08.031.

Riveiro, Alba Redó, and Joshua Mark Brickman. 2020. ‘From Pluripotency to Totipotency: An Experimentalist’s Guide to Cellular Potency’. Development 147 (16). https://doi.org/10.1242/dev.189845.

Roccio, Marta, Daniel Schmitter, Marlen Knobloch, Yuya Okawa, Daniel Sage, and Matthias P. Lutolf. 2013. ‘Predicting Stem Cell Fate Changes by Differential Cell Cycle Progression Patterns’. Development 140 (2): 459–70. https://doi.org/10.1242/dev.086215.

Rothová, Michaela Mrugala, Alexander Valentin Nielsen, Martin Proks, Yan Fung Wong, Alba Redó Riveiro, Madeleine Linneberg-Agerholm, David Eyal, Ido Amit, Ala Trusina, and Joshua Mark Brickman. 2022. ‘Identification of the Central Intermediate in the Extra-Embryonic to Embryonic Endoderm Transition through Single Cell Transcriptomics’. Nature Cell Biology in press.

Saiz, Néstor, Kiah M. Williams, Venkatraman E. Seshan, and Anna-Katerina Hadjantonakis. 2016. ‘Asynchronous Fate Decisions by Single Cells Collectively Ensure Consistent Lineage Composition in the Mouse Blastocyst’. Nature Communications 7 (1): 13463. https://doi.org/10.1038/ncomms13463.

Sakaue-Sawano, Asako, Hiroshi Kurokawa, Toshifumi Morimura, Aki Hanyu, Hiroshi Hama, Hatsuki Osawa, Saori Kashiwagi, et al. 2008. ‘Visualizing Spatiotemporal Dynamics of Multicellular Cell-Cycle Progression’. Cell 132 (3): 487–98. https://doi.org/10.1016/j.cell.2007.12.033.

Sandler, Oded, Sivan Pearl Mizrahi, Noga Weiss, Oded Agam, Itamar Simon, and Nathalie Q. Balaban. 2015. ‘Lineage Correlations of Single Cell Division Time as a Probe of Cell-Cycle Dynamics’. Nature 519 (7544): 468–71. https://doi.org/10.1038/nature14318.

Singer, Zakary S., John Yong, Julia Tischler, Jamie A. Hackett, Alphan Altinok, M. Azim Surani, Long Cai, and Michael B. Elowitz. 2014. ‘Dynamic Heterogeneity and DNA Methylation in Embryonic Stem Cells’. Molecular Cell 55 (2): 319–31. https://doi.org/10.1016/j.molcel.2014.06.029.

Strebinger, Daniel, Cédric Deluz, Elias T Friman, Subashika Govindan, Andrea B Alber, and David M Suter. 2019. ‘ Endogenous Fluctuations of OCT 4 and SOX 2 Bias Pluripotent Cell Fate Decisions ‘. Molecular Systems Biology 15 (9): 1–19. https://doi.org/10.15252/msb.20199002.

Suemori, Hirofumi, Yuzo Kadodawa, Norio Nakatsuji, Koji Goto, Isato Araki, and Hisato Kondoh. 1990. ‘A Mouse Embryonic Stem Cell Line Showing Pluripotency of Differentiation in Early Embryos and Ubiquitous β-Galactosidase Expression’. Cell Differentiation and Development 29 (3): 181–86. https://doi.org/10.1016/0922-3371(90)90120-L.

Waisman, Ariel, Federico Sevlever, Martín Elías Costa, María Soledad Cosentino, Santiago G. Miriuka, Alejandra C. Ventura, and Alejandra S. Guberman. 2019. ‘Cell Cycle Dynamics of Mouse Embryonic Stem Cells in the Ground State and during Transition to Formative Pluripotency’. Scientific Reports 9 (1): 1–10. https://doi.org/10.1038/s41598-019-44537-0.

Wianny, F., F.x. Real, C.l. Mummery, M. Van Rooijen, J. Lahti, J. Samarut, and P. Savatier. 1998. ‘G1-Phase Regulators, Cyclin D1, Cyclin D2, and Cyclin D3: Up-Regulation at Gastrulation and Dynamic Expression during Neurulation’. Developmental Dynamics 212 (1): 49–62.

Yamamoto, Takuya, Miki Ebisuya, Fumito Ashida, Kazuo Okamoto, Shin Yonehara, and Eisuke Nishida. 2006. ‘Continuous ERK Activation Downregulates Antiproliferative Genes throughout G1 Phase to Allow Cell-Cycle Progression’. Current Biology 16 (12): 1171–82. https://doi.org/10.1016/j.cub.2006.04.044.

Yamanaka, Y., F. Lanner, and J. Rossant. 2010. ‘FGF Signal-Dependent Segregation of Primitive Endoderm and Epiblast in the Mouse Blastocyst’. Development 137 (5): 715–24. https://doi.org/10.1242/dev.043471.

Ying, Qi Long, Jason Wray, Jennifer Nichols, Laura Batlle-Morera, Bradley Doble, James Woodgett, Philip Cohen, and Austin Smith. 2008. ‘The Ground State of Embryonic Stem Cell Self-Renewal’. Nature 453 (7194): 519–23. https://doi.org/10.1038/nature06968.

